# miR155 triplicated in Down syndrome regulates hippocampal GABAergic neurogenesis in Alzheimer’s disease

**DOI:** 10.1101/2025.09.28.679053

**Authors:** Xiaodong Zhu, Jean-Vianney Haure-Mirande, Mesude Bicak, Pengfei Dong, Ilya Kruglikov, Aisha Al-subaie, Valentina Fossati, Scott Noggle, Sam Gandy, Michelle E. Ehrlich

## Abstract

MicroRNA dysregulation is implicated in neurodegenerative disorders, including Alzheimer’s disease (AD). The role of neuronal microRNA155 (miR155), elevated in both AD and Down syndrome (DS), remains unknown. We found that *MIR155HG* (miR155 host gene) colocalizes with *APP* (amyloid-beta precursor protein) in a neuron-specific, topologically-associated domain (TAD) within a regulatory network, in the obligate portion of chromosome 21 triplicated in DS which causes AD neuropathology and in most cases, dementia. We investigated miR155 role during neuron development and then validated these findings in an amyloidopathy model. In human induced pluripotent stem cell (hiPSC)-derived neural stem cells (NSCs), cortical neurons and cortical organoids, *MIR155* deletion enhanced NSC proliferation, ventral patterning and GABAergic interneuron generation. However, *MIR155* overexpression inhibited NSC marker expression and GABAergic interneuron generation. *MIR155* upregulates its mRNA targets, *NR2F1*/*2*, key modulators of hippocampal GABAergic interneuron development. In an amyloidopathy mouse model, *miR155* deletion induced the expansion of hippocampal NSCs and increased hippocampal GABAergic interneurons. These findings reveal previously unrecognized miR155 roles in NSC dynamics and GABAergic interneuron development which directionally diverge from extensively studied microglial miR155 in their beneficial vs negative impact on AD mouse models, suggesting that approaching miR155 therapeutically may require balancing the effects in neurons and microglia.

## Introduction

MicroRNAs (miRNAs) are short, 21 to 23 nucleotide, single-stranded, non-coding RNA molecules. They bind most commonly to the 3’UTR (Untranslated Region) of their target mRNAs and repress protein production by destabilizing the mRNA, cleavage of the mRNA strand, and translational silencing (Bartel 2004). In mammalian brains, miRNA expression profiles are cell type-specific (He et al. 2012; Nowakowski et al. 2018), with distinct functions and roles across different neural subtypes, influencing both brain development and susceptibility to neurological disorders (Im and Kenny 2012; Salta and De Strooper 2017). *Mir155* is a multifunctional miRNA first described as a central player in the macrophage inflammatory response (O’Connell et al. 2007) and then extensively studied as a microglial master regulator in neuroinflammation in neurodegenerative conditions, including Alzheimer’s disease (AD) (Butovsky and Weiner 2018; Yin et al. 2023).

Recent studies, including from our laboratory, demonstrated that *miR155* is upregulated in the hippocampal neurons of AD patients and in mouse models of AD pathology (Readhead et al. 2018; Sierksma et al. 2018; Readhead et al. 2020). Other studies have focused on the increase of *miR155* as part of Trisomy 21 in DS patients (Bras et al. 2018) showing impaired neurogenesis and synaptogenesis during human brain development (Russo et al. 2024), that is also manifested in cortical neurons derived from DS patient hiPSCs (Lu et al. 2013; Ovchinnikov et al. 2018), and in the Ts65Dn mouse model of DS (Reeves et al. 1995). The hippocampus plays an essential role in human cognition and memory and is severely affected in AD (Braak and Braak 1990). Adult hippocampal neurogenesis (AHN) (Eriksson et al. 1998; Lee and Thuret 2018) contributes to hippocampal plasticity through the integration of new neurons into existing circuits (Goncalves et al. 2016). AHN in rodents (Kempermann et al. 2004; Goncalves et al. 2016) involves radial glia-like neural stem cells (RGL-NSCs) in the SGZ of the hippocampal dentate gyrus (DG), which generate proliferating intermediate progenitor cells (IPCs) and neuroblasts that differentiate into dentate granule neurons. Markers of hippocampal neurogenesis decline sharply in AD (Moreno-Jimenez et al. 2019; Tobin et al. 2019; Zhou et al. 2022). Hippocampal GABA(gamma-aminobutyric acid)ergic interneurons primarily originate from the ventral telencephalon, and their generation and specification are governed by transcription factors such as NR2F1/2 (Pelkey et al. 2017; Wamsley and Fishell 2017). They represent 10-15% of the total neuronal population and serve as major determinants of virtually all aspects of cortical circuit function (Pelkey et al. 2017). Parvalbumin (PV^+^) and somatostatin (SST^+^) subtypes comprise ∼70% of inhibitory GABAergic interneurons (Pelkey et al. 2017; Wamsley and Fishell 2017). In AD, there are significant reductions in levels of GABA and somatostatin in the cerebrospinal fluid (CSF) and brain (Bareggi et al. 1982; Grouselle et al. 1998; Palop and Mucke 2016). Single-cell RNA sequencing studies confirmed the selective depletion of PV- and SST-positive GABAergic inhibitory neurons (Cain et al. 2023; Mathys et al. 2023; Gabitto et al. 2024).

To determine if miR155 overexpression individually contributes to the changes described above, we utilized 1) *in silico* analyses of published databases to show the relationship in three-dimensional chromatin structure between transcription of *MIR155HG* and *APP*; 2) newly generated *MIR155*-deleted hiPSCs using CRISPR/Cas9 genome editing technology (Jinek et al. 2012; Cong et al. 2013) to produce NSCs, cortical and GABAergic neurons and cortical organoids; 3) lentivirus-mediated *MIR155*-overexpressing hiPSC-derived NSCs, cortical and GABAergic neurons; and 4) *miR155* KO mice to compare and contrast effects of decreased *miR155 in vitro* and *in vivo*. Integration of data from these systems supports a novel role for *MIR155* in regulation of interneuron numbers and phenotypes highly relevant to AD and DS, likely dependent on NR2F1 and NR2F2. Taken together, our findings suggest distinct cell-type-specific roles of *miR155* in AHN and GABAergic interneurons in the pathogenesis of AD. This raises the possibility that therapeutic modulation of *miR155* in both neurons and microglia might be beneficial in the treatment and/or prevention of AD, but the cell-type-specific valence creates a unique challenge, as altering *miR155* expression in either direction or at different stages of disease could be harmful in one cell type while protective in another (Gandy and Ehrlich 2023).

## Results

### Colocalization of APP and MIR155HG genes in a neuron-specific chromatin TAD within a regulatory network on human chromosome 21

*MIR155* is encoded by the host gene, *MIR155HG*, also known as the B-cell Integration Cluster (*BIC*) gene. It is composed of 3 exons that span a 13 kb region on human chromosome 21 (Thai et al. 2007). *MIR155HG* is in the obligate Down syndrome locus, proximal to the *APP* gene which contains 18 exons spanning 290 kb (Supplemental Fig. S1A). Both are overexpressed in Down syndrome patients, all of whom develop AD neuropathology (Bras et al. 2018; Russo et al. 2024). We analyzed genome-wide maps of chromatin accessibility, ATAC-sequencing (assay for transposase-accessible chromatin with high throughput sequencing) data from neuronal nuclei isolated from the superior temporal gyrus (STG) and entorhinal cortex (EC) of AD cases and controls from the Mount Sinai Brain Bank AD (MSBB-AD) cohort (Bendl et al. 2022). In addition, we analyzed H3K4me3 (H3-lysine 4 trimethylation) and H3K27ac (H3-lysine 27 acetylation) ChIP-sequencing (chromatin immunoprecipitation sequencing) data from prefrontal cortex (PFC) neurons of adult control brains (Girdhar et al. 2022). Within *MIR155HG*, we identified ATAC-sequencing peaks in exon 3, H3K4me3 ChIP-sequencing peaks in the transcription start site and exon 3, and H3K27ac ChIP-sequencing peaks in exon 3 (Fig. 1A). These histone markers and chromatin accessibility strongly suggest that *MIR155HG* and *APP,* along with other genes proximal to *APP* that are associated with AD(e.g., *MRPL39*, *JAM2* and *GABPA*; Supplemental Fig. S1A) (Nazarian et al. 2019), are actively transcribed in human neurons (Fig. 1A).

**Figure 1.**
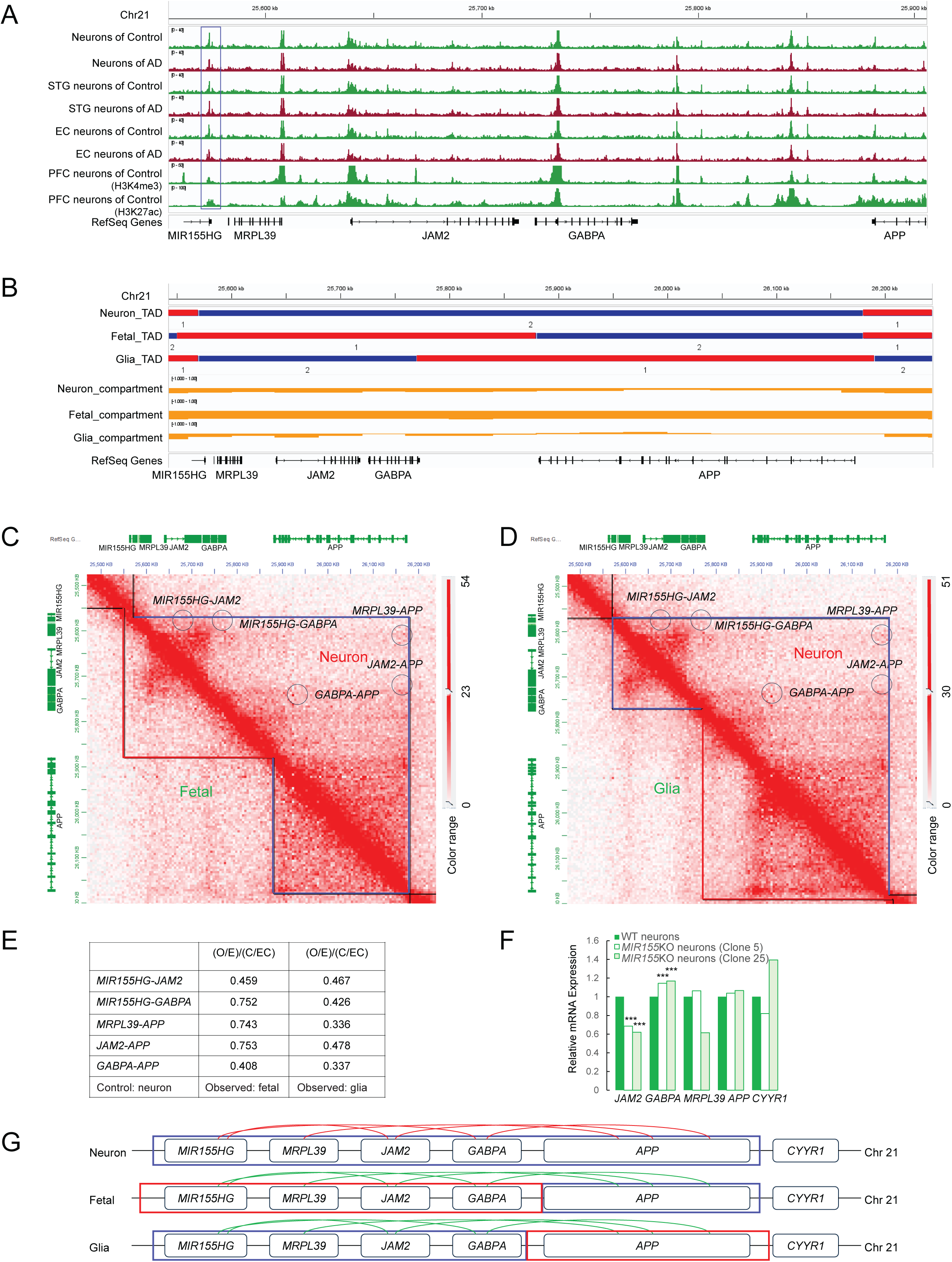
Colocalization of *APP* and *MIR155HG* genes in a neuron-specific chromatin TAD within a regulatory network. (A) ATAC-seq peaks in STG and EC neurons isolated from AD patients and controls, H3K4me3 and H3K27ac ChIP-seq peaks in PFC neurons of adult control brains visualized by using the Integrated Genome Viewer (IGV). The peaks in exon 3 of *MIR155HG* are marked with a blue rectangle. (B) The *APP* gene-centered TADs and compartments on chromosome 21 were identified in neuronal, fetal and glial nuclei and were visualized using the IGV. Individual TADs were labeled with blue (“2”) or red lines (“1”). Changes in compartment scores were demonstrated with the orange line. (C, D) Chromatin contact matrices from the 3D genome browser JuiceBox comparing merged Hi-C sequencing data in adult neurons (NeuN ^+^ cells, top-right triangles in (C) and (D)) versus fetal cortical plate tissue (C, bottom-left triangle) and adult neurons versus glia (NeuN-cells) (D, bottom-left triangle) for a 0.7-Mb locus (chr21:25,500,000-26,200,000) centered on *APP* gene at 5 kb resolution. The Hi-C color scale ranges normalized contact frequencies within this locus. Compared to fetal (C) and glia (D), the neuron-specific chromatin contacts were marked with circles. Top-right blue right angles in (C) and (D) spanning the genomic region between *MIR155HG* and *APP* genes represent a neuron-specific chromatin TAD, however in fetal (C) or glial nuclei (D), the corresponding genomic region was divided into two chromatin TADs which are further marked with a red or blue right angle on the bottom-left of the figure. (E) Comparative analysis (neuron vs fetal and neuron vs glia) of the frequency of chromatin interacting regions of *MIR155HG*-*JAM2, MIR155HG-GABPA, MRPL39-APP*, *JAM2-APP* and *GABPA*-*APP* were summarized in the table. Observed value (O), Expected value (E), Control value (C), Expected Control value (EC), (O/E)/(C/EC) = (Observed value/Expected value)/(Control value/Expected Control value). Compared to Control (neuron), the interaction frequency (Observed, fetal or glia) significantly decreased. (F) Dynamic expression profiles of *JAM2, GABPA, MRPL39, APP* and *CYYR1* in cortical neurons derived from wild type (WT), *MIR155*KO (Clone 5) or *MIR155*KO (Clone 25) hiPSCs, which were generated from RNA sequencing datasets (seen in Supplemental Figs. S1-5, Supplemental table S4), are illustrated as a chart. *** adjusted p value (adj.P.Val) < 0.001. (G) Schematic illustration of localization of *APP* and *MIR155HG* genes in chromatin TADs within regulatory networks. Here chromatin TADs in neuronal (top), fetal (middle) and glial nuclei (bottom) were shown. Blue or red rectangles represent the transition of individual chromatin TADs. Red and green curves indicate the strong and weak interactions among DNA fragments of different genes, respectively.

Chromosomes are partitioned into megabase-sized compartments, named A and B, which contain active and repressed chromatin, respectively. Compartments are further separated into self-interacting neighborhoods called topologically-associated domains (TADs), the disruption of which plays an important role in transcriptional alterations in diseases (Lieberman-Aiden et al. 2009; Dixon et al. 2012). The published data of *in situ* Hi-C sequencing, a method to comprehensively detect genome-wide chromatin interactions in the nucleus, were generated from nuclei isolated from human fetal cortical plate tissue from 18–24 weeks post-conception (referred as “fetal”) and from neurons and glia sorted from adult prefrontal cortex (referred as “neuron” and “glia”) (Hu et al. 2021; Bendl et al. 2022; Rahman et al. 2023). Compared to those in fetal nuclei, there were dramatic transitions in chromosome compartments around the *APP* gene in neurons and glia (Fig. 1B). We found a neuron-specific, *APP*-centered TAD which included genes proximal to *APP*, including *MIR155HG, MRPL39*, *JAM2* and *GABPA* (Fig. 1B). Moreover, compared to those in fetal or glial nuclei, there were significant increases in the frequency of chromatin interacting regions in this neuron-specific TAD, particularly in the interacting regions of *MIR155HG*-*JAM2, MIR155HG-GABPA, MRPL39-APP*, *JAM2-APP* and *GABPA*-*APP* (Fig. 1C-E). These findings were further supported by *MIR155*KO RNA sequencing data (Supplemental Figs. S1-5 and Fig. 1F), which showed that *MIR155* deletion caused statistically significant changes in the expression of *JAM2 and GABPA*, and by an independent study showing the regulatory impact of the *APP* gene on the expression of proximal genes on chromosome 21 in hiPSC-derived DS neurons (Ovchinnikov et al. 2018). Taken together, these results showed the colocalization of *APP* and *MIR155HG* genes in a neuron-specific chromatin TAD within a regulatory network (Fig. 1G).

### MIR155 is a repressor of proliferation in hiPSC-derived neural stem cells

Mature *MIR155* is processed from a long noncoding primary transcript derived from exon 3 (Thai et al. 2007) (Supplemental Fig. S1B). To obtain insights into the physiological function of *MIR155* in neurons and their precursors, we generated isogenic *MIR155* knockout (*MIR155*KO) hiPSC lines using CRISPR/Cas9 methodology (Jinek et al. 2012; Cong et al. 2013). The cell lines were individually monoclonalized and validated by PCR, DNA electrophoresis, and Sanger sequencing (Supplemental Fig. S1C-E). Further characterization of *MIR155*KO clones by immunostaining, qPCR, and G-band karyotype analysis showed homogeneous expression of the pluripotency markers OCT4, NANOG, SSEA4, and TRA-1-60, absence of random differentiation, and normal karyotypes (Supplemental Fig. S1F-H).

To characterize neural phenotypes caused by *MIR155* deletion, *MIR155*KO hiPSCs, along with wild-type (WT) lines, were differentiated into human NSCs and cortical neurons (Chambers et al. 2009; Qi et al. 2017) (Supplemental Fig. S2A). hiPSC-derived NSCs expressed general markers of NSCs with dorsal forebrain fate including FOXG1, NESTIN, PAX6, SOX1, SOX2 and Ki67 (Fig. 2A). Cortical neurons (TUJ1-positive) expressed TBR1, a marker of layer VI of the developing human cortex, and VGLUT1, a marker of glutamatergic excitatory neurons (Supplemental Fig. S2C). During the differentiation of human NSCs into cortical neurons, qPCR assays of miRNAs revealed that *MIR155-5P* expression peaked in NSCs and dropped dramatically in mature neurons (Fig. 2B), whereas *MIR155-3P* was undetected. In parallel, the expression of neuron-enriched *MIR128*, one of the most abundant miRNAs in the adult mouse and human brain (He et al. 2012; Tan et al. 2013), increased gradually from NSCs to terminally differentiated neurons (Fig. 2B). Our miRNA qPCR results are consistent with published data that revealed enriched *MIR155-5P* expression in maturing deep layer neurons and newborn upper layer neurons in human brain (Nowakowski et al. 2018). *MIR155* deletion promoted the expansion of NSCs, by doubling their proliferation (Fig. 2C). In addition, *MIR155*-deleted NSC clones were less cohesive and included more glial-like progeny than WT. We performed RNA sequencing on samples from WT, *MIR155*KO Clone 5 and *MIR155*KO Clone 25 hiPSC-derived NSCs and mature neurons (Supplemental Fig. S3A-B and Supplemental table S4). As shown in gene set enrichment analysis (GSEA, Fig. 2D-E and Supplemental Fig. S3C-D) and the heatmap (Supplemental Fig. S4A), *MIR155*-deleted NSCs upregulated genes associated with GLIOBLASTOMA_PRONEURAL and ASTROCYTE features (Verhaak et al. 2010; Zhong et al. 2018). This suggests that *MIR155*-deleted human NSCs acquired the identity of RGL-NSCs, including high self-renewal ability, increased proliferation, multilineage potency, and migration capacity. These findings also revealed the critical link between transcriptomic dysregulation and the significant increase in NSC proliferation caused by *MIR155* deletion. Furthermore, *MIR155* deletion led to downregulation of genes related to focal adhesion, extracellular matrix, and cytoskeleton (Supplemental Fig. S3F, S4B-C).

**Figure 2.**
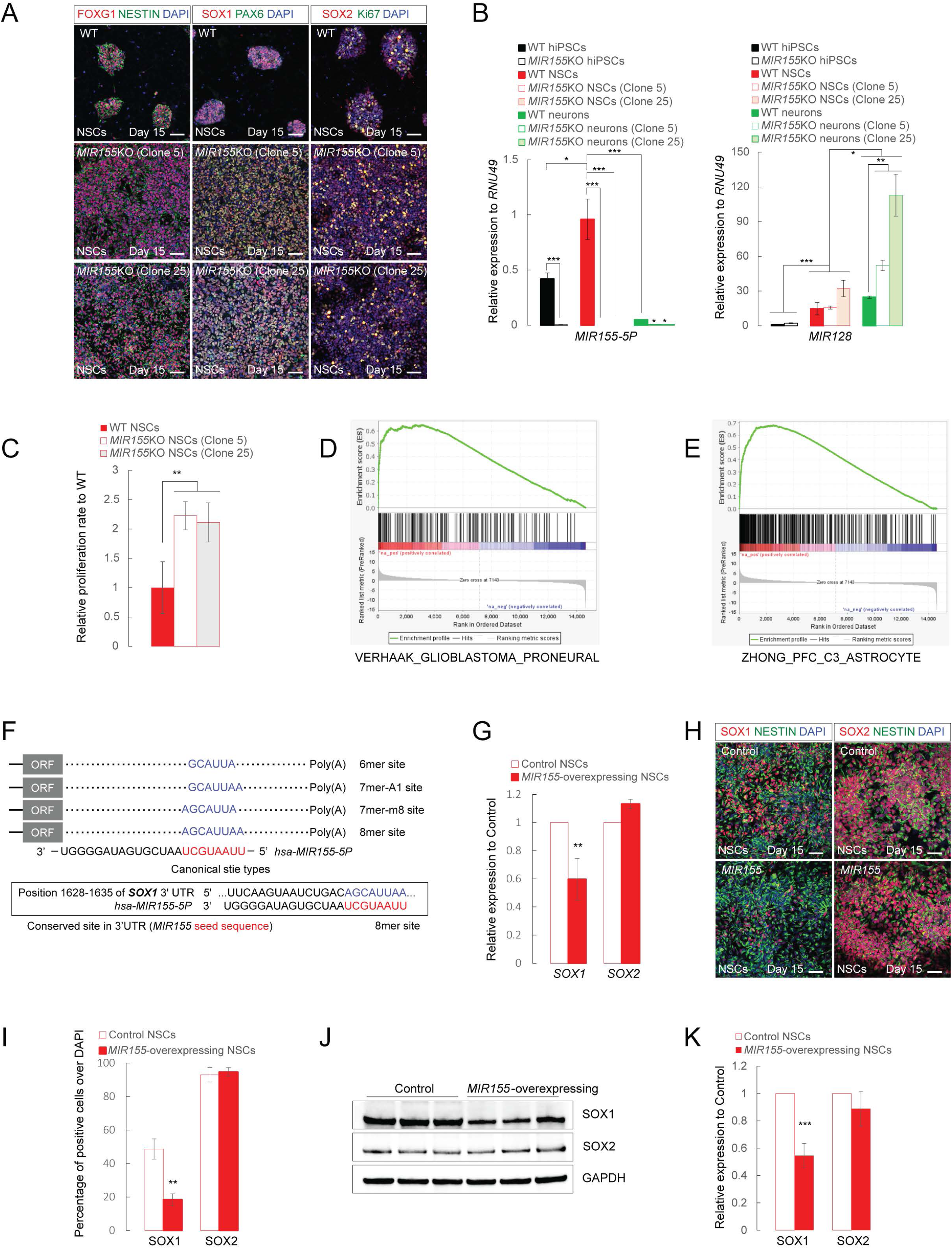
*MIR155* is a repressor of proliferation in hiPSC-derived NSCs. (A) Representative images of wild type (WT), *MIR155*KO Clone 5 and Clone 25 hiPSC-derived NSCs in cell culture at differentiation day 15. In the study, two isogenic *MIR155*KO hiPSC clones, clone 5 and clone 25, were used in parallel with WT in all experiments. Scale bar, 50 µm. (B) The expression levels of *MIR155-5P*, *MIR155-3P* and *MIR128* in hiPSCs, hiPSC-derived NSCs and cortical neurons (in the absence or presence of *MIR155*) were determined by miRNA qPCR. (C) NSCs were seeded at identical densities in 8-well slide chambers and cultured for four days and then labeled with markers of NSCs (A) and DAPI. The 8-well slides were scanned to determine NSC numbers by Agilent BioTek Cytation (4 well replicates of WT, *MIR155*KO Clone 5 or Clone 25 hiPSC-derived NSC culture per each experiment for a total of 3 independent experiments). Y axis refers to relative proliferation speed normalized to WT. **p < 0.01, t test. (D, E) GSEA enrichment plots depicting VERHAAK_GLIOBLASTOMA_PRONEURAL (D) and ZHONG_PFC_C3_ASTROCYTE (E)-associated transcriptional changes in *MIR155*-deleted hiPSC-derived NSCs. Adj.P.Val of DEGs for GESA < 0.05. (F) Canonical miRNA complementary sites. The seed region of *miR155* is marked in red, and 6-nt (6mer), 7-nt (7mer-A1 and 7mer-m8) and 8-nt (8mer) in 3’UTRs of target mRNAs that match to the seed region of *miR155* are marked in blue. The bottom panel shows the conserved 8mer site in 3’UTR of human *SOX1*. (G) The expression levels of *SOX1* and *SOX2* in control and *MIR155*-overexpressing NSCs were determined by qPCR. (H) Representative images of control and *MIR155*-overexpressing NSCs costained for NESTIN (green) and SOX1 (red) or SOX2 (red), along with nuclear counterstaining with DAPI. Scale bar, 50 µm. (I) The percentage of SOX1- or SOX2-positive cells out of total cells (DAPI-positive) in control and *MIR155*-overexpressing NSCs was determined by cell counting. Data represent the mean, n=9 (9 independent biological samples from 3 independent cell cultures). Error bars indicate STDEV. **p<0.01; t test. (J) Representative western blots for SOX1, SOX2 and GAPDH using lysates of control and *MIR155*-overexpressing NSCs. (K) Ratios of SOX1 or SOX2 to GAPDH from (J) showing a significantly lower SOX1 protein level in *MIR155*-overexpressing NSCs than in control NSCs. Data are reported as the mean. n=9 (9 independent biological samples from 3 independent experiments). Error bars indicate STDEV. ***p<0.001; t test. In the qPCR histogram, the y axis indicates relative miRNA level normalized to *RNU49* (B) or relative gene expression normalized to control (I). Data represent mean. Error bars indicate STDEV. Each data point represents nine technical replicates from three independent experiments. *p < 0.05, **p < 0.01 and ***p < 0.001; t test.

To mimic *miR155* upregulation in hippocampal neurons of AD and DS patients and mouse models (Bras et al. 2018; Sierksma et al. 2018; Readhead et al. 2020), we established and characterized lentivirus-mediated *MIR155*-overexpressing human NSCs (Supplemental Fig. S2D-F). SOX1 is an NSC marker (Venere et al. 2012) and three independent programs that invoke evolutionary conservation, i.e. TargetScan (McGeary et al. 2019), miRDB (Liu and Wang 2019) and PicTar (Krek et al. 2005), predict that its mRNA is a *miR155* target due to a conserved octameric site complementary to *MIR155* seed sequence in the 3’UTR of *SOX1* (Fig. 2F). Furthermore, *MIR155* binding to the 3’UTR of *SOX1* is found in a database of experimentally validated mRNA targets of *miR155* (Huang et al. 2022). We found that the SOX1 transcript and protein levels are downregulated in *MIR155*-overexpressing NSCs by about 2-fold (Fig. 2G-K). RNA sequencing on *MIR155*-overexpressing human NSCs and cortical neurons (Supplemental Fig. S7A-B and Supplemental table S5) showed *SOX1* was among the top 5 downregulated DEGs (differentially expressed genes) (Supplemental Fig. S7C), further confirming that *SOX1* mRNA is a target of *MIR155. MIR155*-overexpression led to a marked upregulation of genes related to focal adhesion, extracellular matrix, and cytoskeletal organization (Supplemental Fig. S4D, S7D), showing an opposite pattern to that observed with *MIR155* deletion (Supplemental Fig. S4E). Lastly, a proliferation deficit has been consistently observed in NSCs differentiated from hiPSCs derived from DS patients where *MIR155HG* gene is overdosed (Hibaoui et al. 2014). Taken together, these data suggest that *MIR155* acts as a negative regulator of NSC proliferation.

### MIR155 is a negative regulator of hiPSC-derived GABAergic interneuron development

Our transcriptomic analysis (Supplemental Fig. S3C-D) highlighted upregulation of GABAergic synaptic gene expression in *MIR155*-deleted hiPSC-derived NSC cultures, revealing a previously unknown link between *MIR155* and inhibitory interneuron development. Gene ontology enrichment analysis (Fig. 3A and Supplemental Fig. S3E) and heatmap visualization (Fig. 3B) demonstrated increased expression of genes encoding GABAergic and glutamatergic synaptic proteins following *MIR155* deletion in hiPSC-derived NSCs. These genes include GABA_A_ receptors (*GABRA1*, *GABRB3*, *GABBR1* and *GABRD*), *SLC6A1* (*GAT1*, GABA transporter 1), glutamate decarboxylases (*GAD1* and *GAD2*) and ionotropic glutamate receptor subunits (*GRIA2*, *GRIK2*, *GRIK3*, *GRIK5* and *GRIN3A*). RNA sequencing analysis showed that *MIR155* deletion also induced changes in the expression of neural transcription factors and genes involved in signaling pathways (NOTCH, WNT, BMP, SHH) (Fig. 3C) crucial for self-renewal, proliferation, and neural differentiation. For example, there is upregulation of the SHH-related gene *PTCH2*, which plays key roles in cell proliferation and induction of ventral patterning during neural development (Ericson et al. 1995; Roessler et al. 1996). Multiple transcription factors controlling neurogenesis, cell commitment, and maturation of cortical GABAergic interneurons, originating from the ventral telencephalon (Wamsley and Fishell 2017), were significantly upregulated, including *NKX2-1*, *LHX6*, *DLX1*, *DLX2*, *OLIG1* and *OLIG2* (Pelkey et al. 2017; Wamsley and Fishell 2017), indicating that *MIR155* may negatively regulate GABAergic interneuron development.

**Figure 3.**
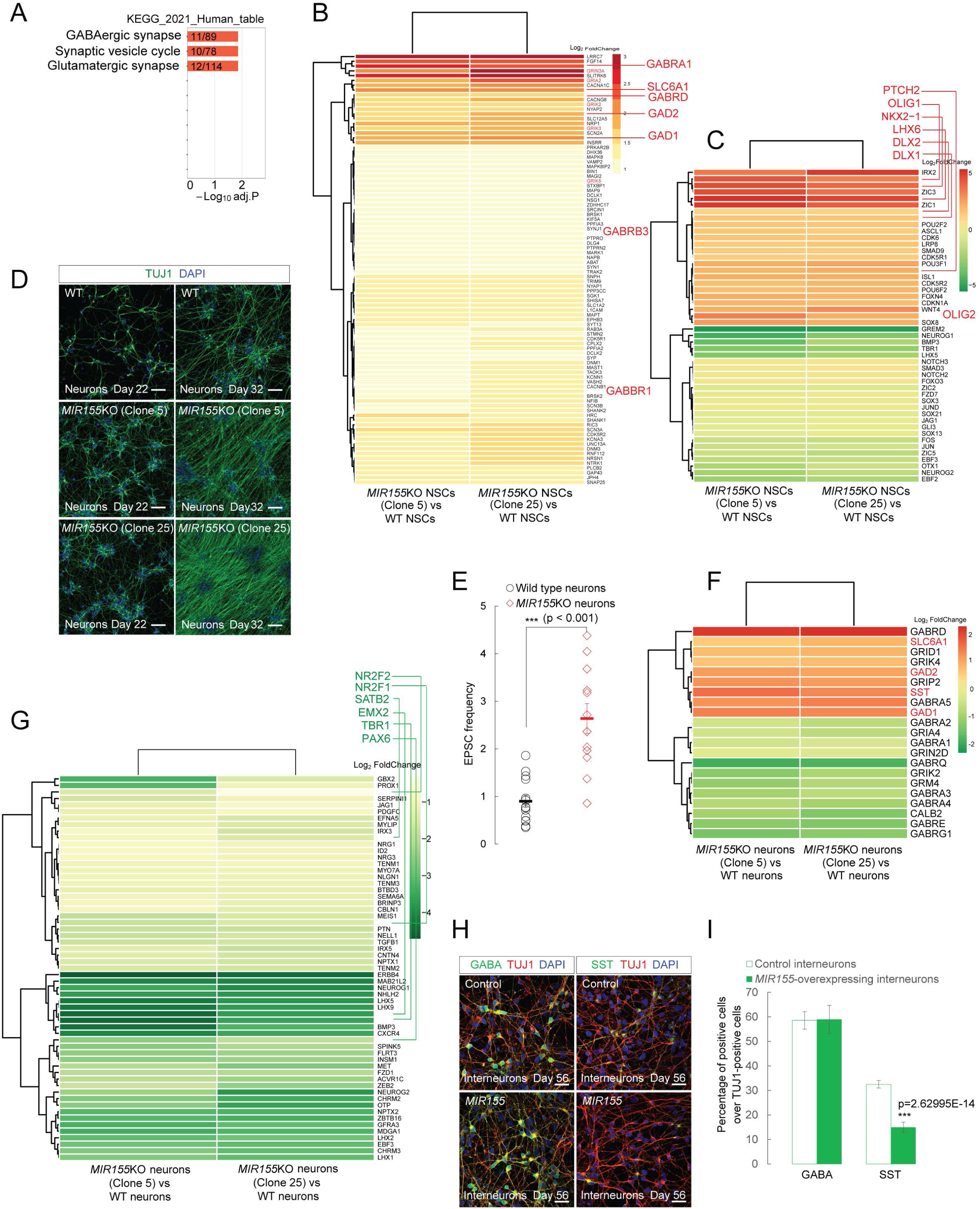
*MIR155* is a developmental regulator of hiPSC-derived GABAergic interneurons. (A) GO terms from differential gene expression analysis of WT and *MIR155*-deficient hiPSC-derived NSCs. Genes upregulated in *MIR155*-deleted NSCs are associated with top statistically significant terms (red histogram) in four modules, here KEGG (Kyoto Encyclopedia of Genes and Genomes) analysis is shown as a bar graph. (B) Heatmap of top statistically significant DEGs encoding GABAergic and glutamatergic synaptic proteins which were upregulated in *MIR155*-deleted NSCs (90 genes derived from GO terms in (Fig. 3A and Supplemental Fig. S3E), typical genes encoding GABA receptors, *SLC6A1*, *GAD1*, *GAD2* and glutamate receptors are marked in red). Adj.P.Val of DEGs shown in the heatmap < 0.05. (C) Expression profiles of neural transcription factors (*NKX2-1*, *LHX6*, *DLX1*, *DLX2*, *OLIG1*, *OLIG2*…) and genes involved in signaling pathways (NOTCH, WNT, BMP, SHH…) are shown as a heatmap. Adj.P.Val of DEGs shown in the heatmap < 0.05. (D) Cortical neurons differentiated from WT, *MIR155*KO Clone 5 and *MIR155*KO Clone 25 hiPSCs were stained for TUJ1 at differentiation day 22 and 32. Scale bar, 50 µm. (E) EPSC (excitatory postsynaptic current) properties were recorded in WT and *MIR155*-deleted hiPSC-derived cortical neurons (n=13 and 12, respectively) at differentiation day 50-60 from two batches of cultures. ***p (=0.000138) < 0.001; t test. (F) Dysregulated expression of GABAergic and glutamatergic signaling components in hiPSC-derived cortical neurons induced by *MIR155* deletion are shown as a heatmap. Adj.P.Val of DEGs shown in the heatmap < 0.05. (G) Heatmap of downregulated genes, dorsal forebrain PAX6-derived cortical neuronal transcription factors including *PAX6*, *TBR1*, *SATB2* and *EMX2* (marked in green), *NR2F1* and *NR2F2* (marked in green). Here 59 genes were derived from the top four GO terms of GO_Cellular_Component in (Supplemental Fig. S5D). Adj.P.Val of DEGs shown in the heatmap < 0.05. (H, I) Control and *MIR155*-overexpressing GABAergic interneurons in culture at day 56 were stained with antibodies against TUJ1 (red, H), GABA (green, H), and markers of GABAergic interneuron subtypes, the calcium-binding protein parvalbumin (PV, undetected) and the neuropeptide somatostatin (SST)(green, H). Scale bar, 50 μm. (I) The percentage of GABA- or SST-positive cells out of total neurons (TUJ1-positive) in control and *MIR155*-overexpressing GABAergic interneurons was determined by cell counting. Data represent the mean, n=9 (9 independent biological samples from 3 independent cell cultures). Error bars indicate STDEV. ***p<0.001; t test.

To further investigate the potential glutamatergic and/or GABAergic phenotype associated with *MIR155*-deletion, we differentiated hiPSCs toward cortical neurons, using a well-established protocol that primarily generates dorsal glutamatergic excitatory neurons, along with a smaller proportion of interneurons (Chambers et al. 2009; Qi et al. 2017) (Supplemental Fig. S2A and S2C). *MIR155*-deleted neurons exhibited enhanced neurite growth (Fig. 3D), consistent with another report (Gaudet et al. 2016). Electrophysiological analysis showed that *MIR155* deletion induced a significant increase in the frequency of spontaneous synaptic events in hiPSC-derived cortical neurons, without affecting neuronal firing properties (Fig. 3E and Supplemental Fig. S2G-H). Transcriptomic analysis (Supplemental Fig. S5A-D) comparing *MIR155*-deleted versus WT neuronal cultures revealed a robust induction of GABAergic interneuron markers, including *GAD1*, *GAD2*, *SLC6A1* and *SST* (Fig. 3F), showing overlap with the genes upregulated in *MIR155*-deleted NSCs, described above. These markers were likely induced at the expense of dorsal forebrain markers such as *EMX2*, *PAX6*, *TBR1* and *SATB2* (Fig. 3G). *MIR155* deletion induced downregulation of *NR2F1* and *NR2F2* (Fig. 3G).

To mimic *miR155* upregulation in hippocampal neurons in AD (Readhead et al. 2020) and DS patients (Bras et al. 2018), we generated *MIR155*-overexpressing hiPSC-derived cortical GABAergic interneurons as described (Maroof et al. 2013) (Supplemental Fig. S2B and S2I-J). Only 37 DEGs were identified in *MIR155*-overexpressing cortical glutamatergic neurons (Supplemental Fig. S2D-F and S7E), raising the possibility that the transcriptomic effects of *MIR155* upregulation may be cell-type specific. We found that *MIR155* overexpression resulted specifically in a significant decrease in the percentage of SST-positive cortical GABAergic interneurons (Fig. 3H-I). These data are consistent with those derived from systems containing the entire triplication (Ross et al. 1984; Kobayashi et al. 1990; Huo et al. 2018), and therefore identify miR155 as a causative gene in DS. GABA receptor genes (*GABRA1*, *GABRA4* and *GABRB2*) and *NR2F2* are included in the list of mRNA targets of *MIR155* (Fig. 4A) according to three independent prediction tools (Krek et al. 2005; Liu and Wang 2019; McGeary et al. 2019). Gene expression analysis by qPCR showed that *MIR155* overexpression induced significant changes in expression of its mRNA targets, downregulating *GABRA1* and *GABRA4*, but upregulating *NR2F2* and NR2F family homolog *NR2F1* (Fig. 4B). Collectively, our studies showed downregulation of *NR2F1* and *NR2F2* expressions in *MIR155*-deleted hiPSC-derived cortical neurons (Fig. 3G), and upregulation of *NR2F1* and *NR2F2* expressions in *MIR155*-overexpressing NSCs (Supplemental Fig. S7C), in *MIR155*-overexpressing cortical neurons (Supplemental Fig. S7E), and in *MIR155*-overexpressing GABAergic interneurons (Fig. 4B). During the transition from NSCs (WT, Control or *MIR155*-overexpressing) to cortical neurons, *MIR155-5P* expression dropped dramatically (Fig. 2B and Supplemental Fig. S2F), and the expression of *NR2F1* and *NR2F2* decreased synchronously (Supplemental Fig. S6A-D, S7F-J and Fig. 4C). A bioinformatics screen using miRBASE identified 106 target genes with the ARE (AU-rich element) motifs in the highly conserved 3’UTRs of their mRNAs complementary to the seed regions of *miR155*, including *NR2F1* and *NR2F2* (Fig. 4D-E). These results support a critical role of *MIR155* in positively regulating the expression of *NR2F1* and *NR2F2*, possibly by stabilizing their mRNAs through *MIR155* binding with the AREs in their 3’UTRs (Vasudevan et al. 2007). In summary, these results point to *MIR155* as an important regulator of the development of GABAergic interneurons or their specific subtypes, and of *NR2F1* and *NR2F2* expressions.

**Figure 4.**
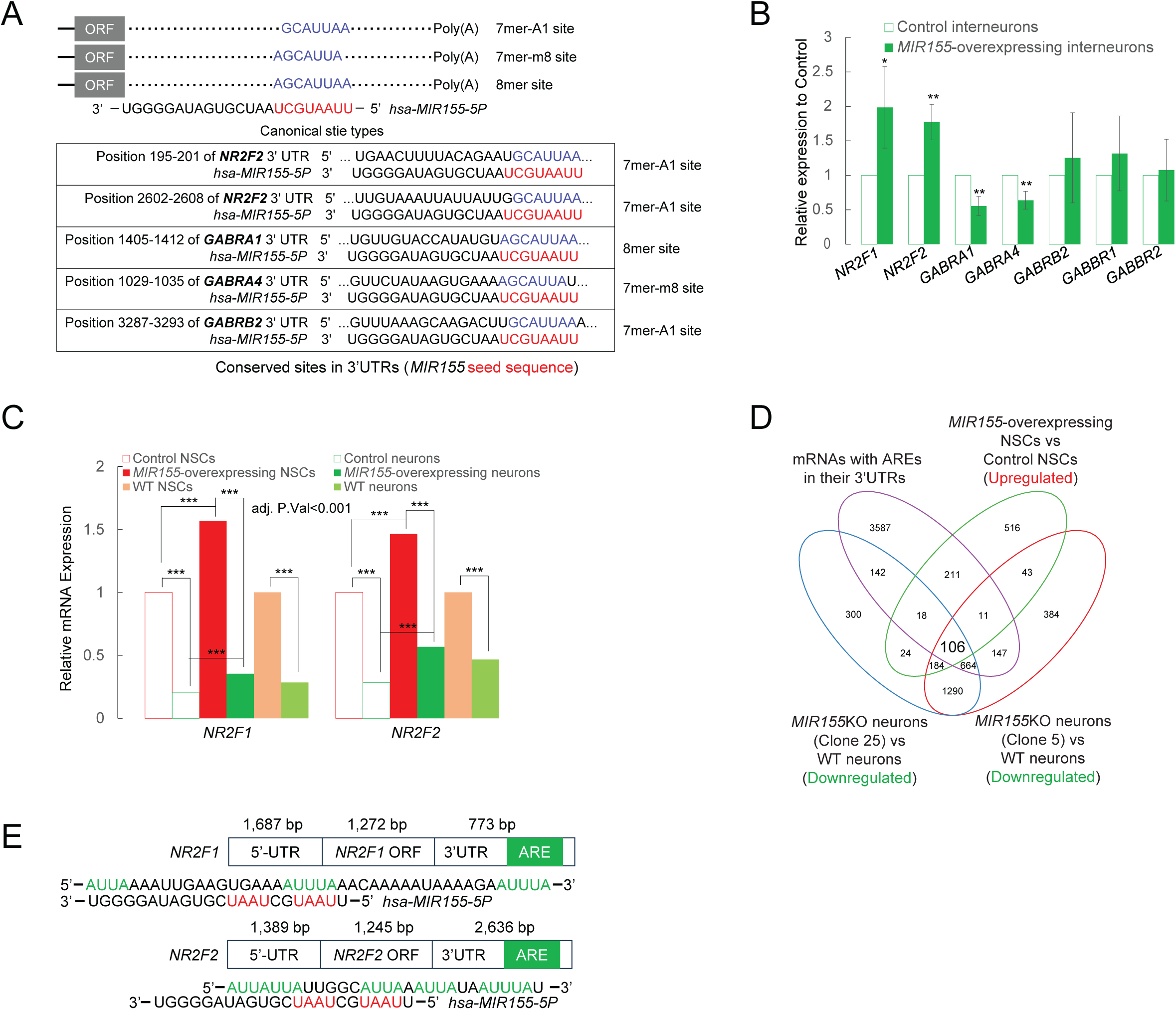
*MIR155* positively regulates expression of its mRNA targets, *NR2F1* and *NR2F2*. (A) Canonical miRNA complementary sites. The seed region of *miR155* is marked in red, and 7-nt (7mer-A1 and 7mer-m8) and 8-nt (8mer) in 3’UTRs of target mRNAs which match to the seed region of *miR155* are marked in blue. The bottom panel shows the conserved sites in 3’UTRs of the mRNA targets of *MIR155*, *NR2F2*, *GABRA1*, *GABRA4* and *GABRB2*. (B) The expression levels of selected genes in control and *MIR155*-overexpressing GABAergic interneurons were determined by qPCR. In the qPCR histogram, the y axis indicates relative gene expression level normalized to control. Data represent mean. Error bars indicate STDEV. Each data point represents 9 technical replicates from 3 independent experiments. *p < 0.05, **p < 0.01; t test. (C) Dynamic expression profiles of *NR2F1* and *NR2F2* during the transition from NSCs (WT, control or *MIR155*-overexpressing) to cortical neurons are illustrated as a chart. ***adj.P.Val < 0.001. (D) Intersections of the downregulated (green) DEGs derived from the comparison of *MIR155*-deleted (Clone 5 or Clone 25) versus WT human cortical neurons, the upregulated (red) DEGs derived from the comparison of *MIR155*-overexpressing versus control NSCs and predicted mRNA targets with AREs in their 3’UTRs, are shown as a Venn diagram. (E) The AREs and ARE motifs (AUUA and AUUUA) in the 3’UTRs of mRNAs of human *NR2F1* and *NR2F2* are marked in green. The seed sequences (complementary to ARE motifs) in *MIR-155-5P* are marked in red.

### MIR155 deletion enhanced the ventral patterning and the induction of GABAergic interneurons in 3D cortical organoids

We next determined whether the phenotypes of *MIR155* deletion in 2D hiPSC-derived cell cultures could be recapitulated in more complex 3D human cortical organoids, prepared using a modified version of an established protocol (Fig. 5A) (Madhavan et al. 2018). We have demonstrated that cortical organoids recapitulate regional organization and cortical cell types present in the developing human brain, when compared to standard cerebral organoids (Lancaster et al. 2013; Kalpana et al. 2024). Cortical organoids at early stages (around 10 days) formed an abundance of large rosette-like neuroepithelia, which expressed SOX1, PAX6, SOX2 and N-Cadherin (Fig. 5B and 5D) and surrounded a fluid-filled cavity resembling a ventricle with characteristic apical localization of the neural specific N-Cadherin (also called neural rosette, Fig. 5D). Compared to the organoids derived from WT hiPSCs, *MIR155*KO organoids contained many more PAX6- and SOX1-positive NSCs (Fig. 5B-C) and the density of SOX2- and N-Cadherin-positive neural rosettes was significantly higher (Fig. 5D-E). The size of neural rosettes in *MIR155*-deleted hiPSC-derived organoids was slightly smaller (Fig. 5D and 5F).

**Figure 5.**
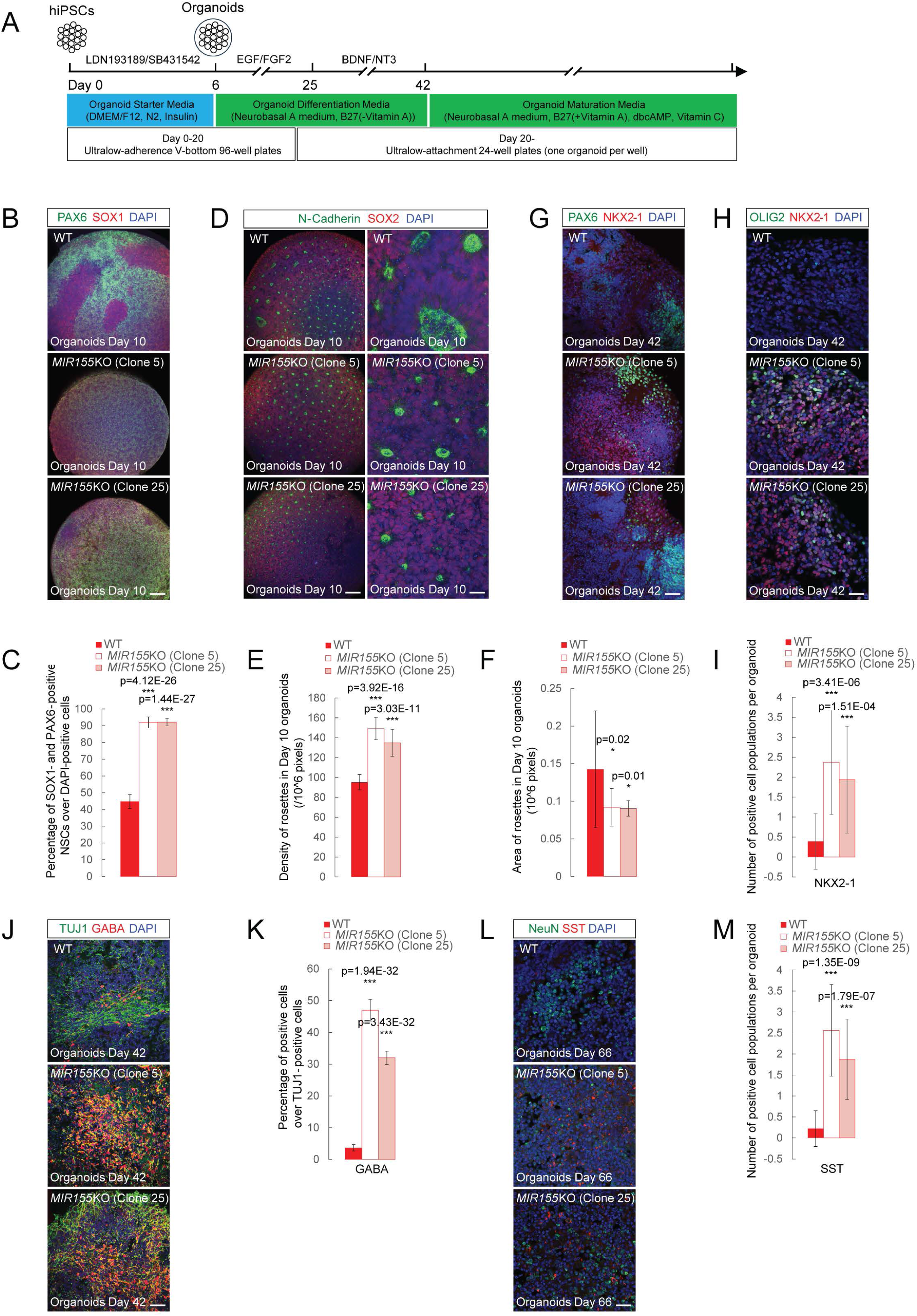
*MIR155* deletion enhanced the ventral patterning and the GABAergic interneuron generation in 3D cortical organoids. (A) Schematic for generating cortical organoids from hiPSCs. The culture conditions, media and cocktails of growth factors or small molecules are indicated. (B, D) 10-day cortical organoids derived from WT, *MIR155*KO Clone 5 or Clone 25 hiPSCs were immunostained with antibodies against SOX1 (red) and PAX6 (green) (B), SOX2 (red) and N-Cadherin (green) (D). Scale bar, 100 µm (B and left panel of D) and 40 µm (right panel of D). (C, E, F) The percentage of PAX6- and SOX1-positive NSCs over total cells (DAPI-positive) (C), the density (E) and area (F) of neural rosettes in 10-day cortical organoids. The numbers of PAX6- and SOX1-positive NSCs, total cells (DAPI-positive) and neural rosettes were counted with ImageJ software. Neural rosettes and the corresponding organoid were outlined by manually tracing and then the area per rosette or organoid was measured with ImageJ software. (G, H, I) Representative images of neural progenitors ((G) NKX2-1 (ventral, red) and PAX6 (dorsal, green)) and GABAergic interneuron progenitors ((H) NKX2-1 (ventral, red) and OLIG2 (ganglionic eminence, green)) in cortical organoids at week 6 of differentiation. Scale bar, 40 µm. (I) The number of NKX2-1-positive cell populations per organoid at week 6 of differentiation. (J, K) Cortical organoids at week 6 of differentiation, which were derived from WT, *MIR155*KO Clone 5 or Clone 25 hiPSCs, were immunostained with antibodies against TUJ1 (green) and GABA (red, J), along with nuclear counterstaining with DAPI. Scale bar, 40 µm. (K) The percentages of GABA-positive cells out of total neurons (TUJ1-positive) in organoids at week 6 of differentiation. (L, M) Representative images of SST-positive GABAergic interneurons (L, co-stained with NeuN) in cortical organoids at day 66 of differentiation. Scale bar, 40 µm. (M) The number of SST-positive cell populations per organoid at day 66 of differentiation. All image analysis of cortical organoids were performed on the images (n = 16) from 16 organoids per batch of organoid culture for a total of 3 independent batches of organoid cultures. *p < 0.05, ***p < 0.001. Data represent the mean, Error bars indicate STDEV.

The organoids ultimately developed into concentric multilayer structures composed of NPCs (neural progenitor cells), lower (CTIP2^+^) and upper (SATB2^+^) cortical layer neurons (data not shown). At 6 weeks, the number of ventral NPCs (NKX2-1^+^) in *MIR155*-deleted hiPSC-derived organoids had increased significantly compared to WT organoids (Fig. 5G and 5I), consistent with the *MIR155*-deletion phenotype observed in 2D hiPSC-derived NSC cultures (Fig. 3C). Given that NKX2-1 is a marker of ventral prosencephalic progenitor populations (Sussel et al. 1999), our results suggest that *MIR155* works as a repressor of ventral patterning during brain development. The NKX2-1 domain in the human embryonic ventral forebrain is further subdivided into an OLIG2-negative preoptic-area anlage, and an OLIG2^+^ ganglionic eminence where cortical and hippocampal GABAergic interneurons arise (Maroof et al. 2013; Pelkey et al. 2017). Thus, to confirm a link between *MIR155* and GABAergic interneuron development, we quantified the number of OLIG2^+^ progenitors and found that nearly 40% of the NKX2-1-positive cells in the organoids co-expressed OLIG2 (Fig. 5G-H). Furthermore, *MIR155*-deleted organoids contained significantly more GABA^+^/TUJ1^+^ interneurons than WT organoids (Fig. 5J-K). In a more mature stage (day 66 of differentiation), cortical organoids derived from WT hiPSCs contained few SST-positive GABAergic interneuron subtype, however, the number of SST-positive GABAergic interneurons increased significantly in *MIR155*-deleted hiPSC-derived organoids (Fig. 5L-M). PV-positive GABAergic interneurons were not detected, likely because of their later development during postnatal and adolescent stages, a time frame difficult to capture in hiPSC-derived models (Hu et al. 2014; Pelkey et al. 2017; Wamsley and Fishell 2017). Taken together, these results indicate that *MIR155* deletion enhanced the ventral patterning and the induction of GABAergic interneurons in cortical organoids.

### Identification of mRNA targets of MIR155 in NSCs and cortical neurons

To identify additional potential targets of *MIR155* and a possible link between gene expression changes in its mRNA targets and developmental regulation of NSCs and GABAergic interneurons, we used three computational miRNA target prediction tools (Krek et al. 2005; Liu and Wang 2019; McGeary et al. 2019) and combined the results with a published database of experimentally validated mRNA targets of *miR155* (Huang et al. 2022) (Supplemental Fig. S8A). Intersection of the gene lists from multiple prediction tools yielded 14 targets in hiPSC-derived NSCs (Supplemental Fig. S8B-D), including *GABRA1*, *TBR1*, a genetic determinant for NSC differentiation (Hevner et al. 2001; Papaioannou 2014), and *FGF14*, a regulator of GABAergic inhibitory synaptic transmission (Di Re et al. 2017), and 69 targets in cortical neurons (Supplemental Fig. S8E-H), including *NR2F2*, *SOX1*, an NSC marker (Venere et al. 2012), *SATB2,* and *PROX1*, a developmental determinant for hippocampal NSCs (Stergiopoulos et al. 2014). Additionally, we highlighted a shortlist of putative candidate genes for further investigation (Supplemental Fig. S8I-J). A significant proportion of well-studied mRNA targets of *MIR155* was downregulated in *MIR155*-deficient hiPSC-derived NSCs and cortical neurons (e.g., *NR2F2*), indicating the involvement of other transcriptional regulators in the expression of DEGs identified in our RNA sequencing datasets. By performing ChIP Enrichment Analysis (ChEA), we observed an involvement of epigenetic regulators in the expression of DEGs, including the SUZ12-JARID2-EZH2 complex (Supplemental Fig. S8K-N), where JARID2, a regulator of neural development (Takeuchi et al. 1995), is a verified mRNA target of *miR155* (Escobar et al. 2014). Loss-of-function mutations in *JARID2* gene cause a neurodevelopmental syndrome, characterized by developmental delay, cognitive impairment and autistic features (Cadieux-Dion et al. 2022). We detected dynamic changes in the expression of several validated mRNA targets of *MIR155*, including *MEIS1*, *CEBPB* and *CLDN1*. These findings indicate *MIR155*-mRNA targets play key roles during the differentiation of NSCs into cortical neurons (Supplemental Fig. S6A-D, S7F-J and S8O-P).

### Deletion of miR155 induced the expansion of RGL-NSCs in the SGZ and the hilus and their ectopic localization in the granule cell layer

Based on our work demonstrating mitigation of parts of the phenotype in *APP^KM670/671NL^/PSEN1^Δexon9^* (*APP/PS1*) (Jankowsky et al. 2004) double transgenic mice (Readhead et al. 2018; Readhead et al. 2020), we hypothesized that *miR155* may have similar functions in adult mouse hippocampal NSCs to those in hiPSC-derived NSCs. Using a miRNAscope assay, we observed a significant enrichment of *miR155* expression in NSCs in the DG and glutamatergic neurons and/or GABAergic interneurons in CA1 (Fig. 6A). In WT mice, the number of Ki67- and Dcx (Doublecortin)-positive cells in the DG declined from 1 to 7 months (Ben Abdallah et al. 2010). In *APP/PS1* mice, hippocampal amyloidosis becomes detectable by 4 months (Radde et al. 2006). We assessed whether *miR155* deletion influences the proliferation of RGL-NSCs. Consistent with *in vitro* results, *APP/PS1-miR155*KO mice showed an increase in Sox1-positive NSCs in both the SGZ and the hilus (Fig. 6B-C) compared to *APP/PS1* mice expressing *miR155*. In 4-month-old *APP/PS1* mice, *miR155* deletion induced an increase in the number of Sox1- and Gfap-positive RGL-NSCs which lined up along the SGZ and were recognized by their characteristic bipolar or unipolar processes (Garcia et al. 2004) (Fig. 6B and 6D). However, there was no change in the overall number of Dcx-positive immature neurons in the SGZ and granule cell layer (GCL) of the DG in hippocampi in 4-month-old *APP/PS1-miR155*KO mice compared with *APP/PS1* mice (Supplemental Fig. S9A-C).

**Figure 6.**
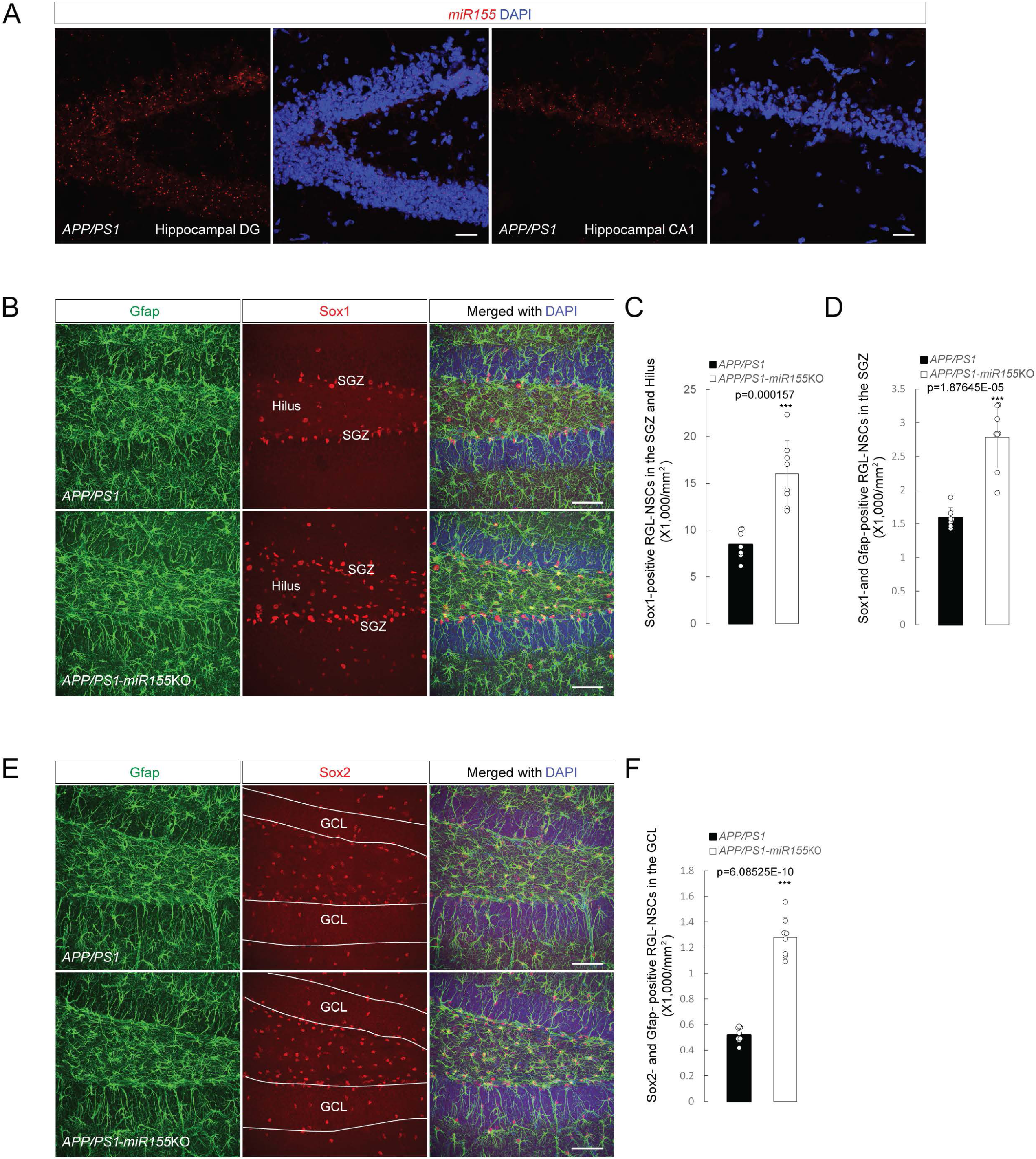
*miR155* deletion induced the expansion of RGL-NSCs in the SGZ and the hilus and their ectopic localization in the GCL. (A) miRNAscope of *miR155* (red) combined with a nuclear counterstain (DAPI) showing *miR155* expression in hippocampal DG and CA1 in 4-month-old *APP/PS1* mice. Scale bar, 20 µm. (B) Representative images of *miR155* deletion-induced expansion in Gfap (green)- and Sox1 (red)-positive RGL-NSCs in the SGZ and hilus of *APP/PS1* mice at 4 months of age. Scale bar, 50 µm. (C) Quantification of total Sox1-positive cells in the SGZ and hilus of *APP/PS1* and *APP/PS1-miR155*KO mice at 4 months of age. (D) Quantification of Sox1- and Gfap-positive cells in the SGZ of 4-month-old *APP/PS1* and *APP/PS1-miR155*KO mice. (E) Representative images of Gfap (green)- and Sox2 (red)-positive RGL-NSCs in the hippocampal GCL of *APP/PS1* and *APP/PS1-miR155*KO mice at 4 months of age showing disruption of *miR155*-induced their ectopic localization in the GCL (marked with white curves). Scale bars, 50 µm. (F) Quantification of ectopically located Gfap- and Sox2-positive cells in the GCL of 4-month-old *APP/PS1* and *APP/PS1-miR155*KO mice. Total numbers of positive cells in the SGZ, the SGZ and hilus, or the GCL were counted in sections from 9 different animals (5 female and 4 male) per group and normalized by the area size. Data presented as mean ± SEM. Error bars indicate STDEV. ***p < 0.001; t test.

Given our *in vitro* observation that *MIR155* regulates genes associated with focal adhesion, extracellular matrix and cytoskeleton (Supplemental Figs. S1-7), we next investigated whether *miR155* deletion would lead to abnormal migration of hippocampal NSCs. While RGL-NSCs were present in an organized fashion along the SGZ in 4-month-old *APP/PS1* mice, *APP/PS1-miR155*KO mice showed more Gfap^+^/Sox2^+^ RGL-NSCs penetrating the GCL, with a 2-fold increase in Gfap^+^/Sox2^+^ RGL-NSCs ectopically located in the GCL (Fig. 6E-F). These results suggest that in *APP/PS1* mice, *miR155* deletion induces the expansion of RGL-NSCs in the SGZ and the hilus as well as their migration into the GCL.

### Deletion of miR155 led to an increase in GABAergic interneuron number in the hippocampi of 8-month-old APP/PS1 mice

Glutamatergic granule cells are the sole neuronal subtype generated from a pool of adult NSCs in the DG of the mature mammalian hippocampus (Kempermann et al. 2004; Goncalves et al. 2016). The integration of these new neurons into functional hippocampal circuits contributes to learning and memory (Kempermann et al. 2004; Goncalves et al. 2016). Our *in vitro* studies with hiPSC cultures indicate that *MIR155* deletion caused induction of GABAergic interneurons (Figs. 3 and 5). Using NeuN and GABA immunofluorescence, we found that in 8-month-old *APP/PS1* mice, *miR155* deletion increased the number of GABA-positive cells in the hippocampal DG, without changing the total number of GABA-positive cells in the whole hippocampus (Fig. 7A-B and Supplemental Fig. S9D-E). Given their ability to control microcircuits and reliably fire at high frequencies, PV neurons are at the center of network synchrony, network oscillations, and memory processing (Sohal et al. 2009; Hu et al. 2014). In AD patients and mouse models, impairments in PV interneurons have been linked to alterations in network hypersynchrony, gamma and theta oscillations, and cognitive deficits (Verret et al. 2012; Iaccarino et al. 2016; Palop and Mucke 2016; Martinez-Losa et al. 2018; Etter et al. 2019; Martorell et al. 2019). In 8-month-old *APP/PS1* mice, *miR155* deletion caused a significant increase in PV^+^ cells in the hippocampi, most prominent in the CA1 region, with no change in the number of Sst^+^ cells (Fig. 7C-D and Supplemental Fig. S9F). These results align well with our previous findings that constitutive absence of *miR155* prevented impaired performance in Barnes maze learning behavior (Readhead et al. 2018; Readhead et al. 2020), implying that PV interneurons are primarily responsible for the beneficial effect. *MIR155* positively regulates its mRNA targets, *NR2F1* and *NR2F2 in vitro* (Fig. 4), and we found that in *APP/PS1* mice, *miR155* deletion significantly downregulated the expression of Nr2f1 (Fig. 7E-H), further confirming miR155 as a positive regulator of expression of Nr2f1.

**Figure 7.**
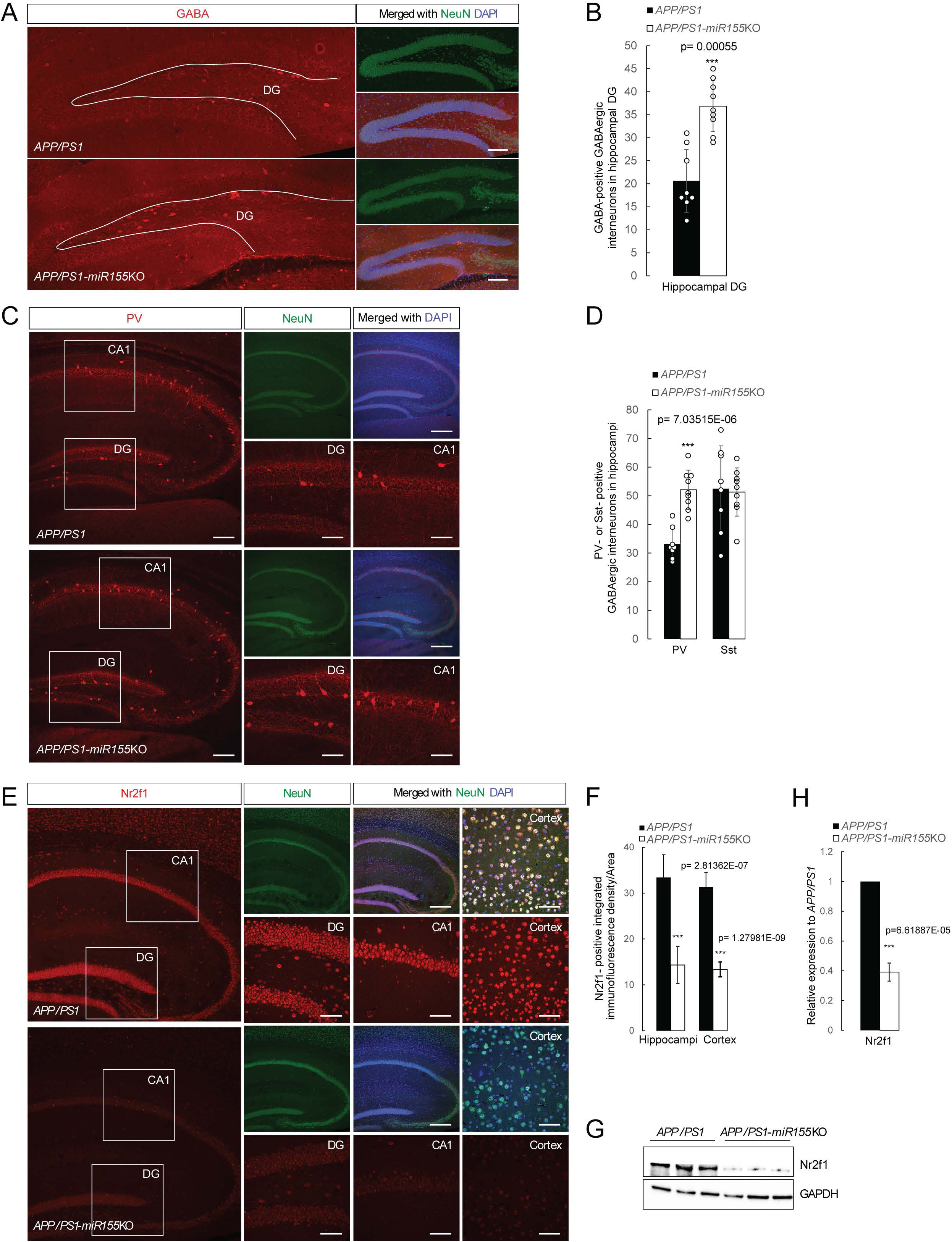
*miR155* deletion caused a significant increase in GABAergic interneurons in hippocampi of 8-month-old *APP/PS1* mice. (A, B) Representative images of hippocampal sections of *APP/PS1* and *APP/PS1-miR155*KO mice at 8 months of age double-immunolabeled for NeuN (green) and GABA (red), along with nuclear counterstaining with DAPI (A). Scale bar, 50 µm. (B) Stereological quantification of GABA-immunoreactive cell number in hippocampal DG (marked with white curves) of 8-month-old *APP/PS1* and *APP/PS1-miR155*KO mice (n=9 mice/group). ***p < 0.001; t test. (C, D) Representative images of hippocampal sections of *APP/PS1* and *APP/PS1-miR155*KO mice at 8 months of age double-immunolabeled for NeuN (green) and parvalbumin (PV)(red, C), along with nuclear counterstaining with DAPI. Scale bar, 100 µm. The images (DG and CA1) on the bottom-right are higher-magnification images from the boxed area on the left. (D) Stereological quantification of PV- or Sst-(Supplemental Fig. S9F) immunoreactive cell numbers in hippocampi of 8-month-old *APP/PS1* and *APP/PS1-miR155*KO mice (n=9 mice/group). ***p < 0.001; t test. (E, F) Representative images of brain sections of 8-month-old *APP/PS1* and *APP/PS1-miR155*KO mice double-immunolabeled for NeuN (green) and Nr2f1 (red, E), along with nuclear counterstaining with DAPI. Scale bar, 100 µm. The two images (DG and CA1) on the bottom-right are higher-magnification images from the boxed area on the left. The neuronal expression of Nr2f1 in cortex is shown in the right higher-magnification images. (F) Quantification of integrated density of Nr2f1 immunofluorescence in brains of 8-month-old *APP/PS1* and *APP/PS1-miR155*KO mice (n=9 mice/group). ***p < 0.001; t test. (G) Representative western blots for Nr2f1 and GAPDH using lysates of the prefrontal cortex of 8-month-old *APP/PS1* and *APP/PS1-miR155*KO mice. (H) The ratio of Nr2f1 to GAPDH from (G) showing a significantly lower Nr2f1 protein level in *APP/PS1-miR155*KO than *APP/PS1* mice. Data are reported as the mean. n=6 mice/group. Error bars indicate STDEV. ***p<0.001; t test.

## Discussion

In this study, we utilized an integrated analysis of published epigenetic datasets, *in vitro* hiPSC technology, and *in vivo* mouse models of AD pathology, to investigate the neuronal expression and function of *MIR155.* We identified a novel, cell-type-specific role for *miR155* in adult hippocampal neurogenesis and GABAergic interneurons within the hippocampus of AD model mice, uniquely demonstrating a well-established link of the Down syndrome gene, *MIR155HG*, to *APP* and AD (Wiseman et al. 2015). We demonstrated that expression of *miR155* is dynamically regulated during the transition of NSCs into terminally differentiated cortical neurons, with a sharp increase in NSCs followed by a decline in mature cortical neurons. These findings are consistent with the notion that expression of *miR155* is developmentally- and cell cycle-regulated in dividing cells, as shown in other systems (Decembrini et al. 2009). Importantly, increased *miR155* expression has been observed in hippocampal neurons in tissues from patients with AD (Sierksma et al. 2018; Readhead et al. 2020) and DS (Bras et al. 2018) or traumatic brain injury (Harrison et al. 2017), and is now also supported by the results of ATAC-seq, H3K4me3 ChIP-seq and H3K27ac ChIP-seq on neuronal nuclei isolated from frozen postmortem human brains. We conclude that *miR155* dysregulation in hippocampal NSCs and GABAergic interneurons contributes to AD pathology through mechanisms independent of miR155 actions in microglia (Yin et al. 2023). Notably, the AD-associated cell-type-specific changes in *miR155* exhibit opposing effects, with the role of microglial *miR155* contrasting with its function in NSCs and GABAergic interneurons.

In both hiPSC-derived and mouse hippocampal NSCs in *APP/PS1* mice (Jankowsky et al. 2004), *miR155* deletion enhanced NSC proliferation, whereas its overexpression downregulated NSC markers. A proliferation deficit has been consistently observed in NSCs differentiated from hiPSCs derived from DS patients (Hibaoui et al. 2014), implying miR155 in Trisomy 21 as a pathogenic gene. Our results are also supported by a study showing that disruption of *miR155* in mice leads to reversal of inflammation-induced decrease in NSC proliferation and neural differentiation in the DG, and that Nestin-specific elevation of *miR155* reduced immature neuron survival and induced ectopic localization of RGL-NSCs in the DG (Woodbury et al. 2015). In *MIR155*-deleted hiPSC-derived NSC cultures, we observed an induction of GABAergic synaptic gene expression. This induction was accompanied by the expression of key transcription factors involved in GABAergic interneuron development, including *NKX2-1*, *LHX6*, *DLX1*, *DLX2*, *OLIG1* and *OLIG2* (Pelkey et al. 2017; Wamsley and Fishell 2017). In cortical organoids, *MIR155* deletion induced a significant increase in the expression of the transcription factor NKX2-1, a molecular determinant of ventral brain development (Sussel et al. 1999). Furthermore, *MIR155* deletion significantly enhanced the generation of GABAergic interneurons in the cortical organoids. These results indicate that *MIR155* acts as a repressive regulator of ventral patterning and of GABAergic interneuron induction during brain development, contrary to the actions of Sonic Hedgehog (Shh) (Ericson et al. 1995; Roessler et al. 1996). The well-validated, commercially available *miR155* inhibitors may be promising tools to induce the ventral patterning and the generation of GABAergic interneurons in brain organoids.

In hiPSC-derived cortical neurons, *MIR155* deletion enhanced neurite growth, caused a robust induction of GABAergic interneurons and downregulated NR2F1 and NR2F2 (Kanatani et al. 2015; Hu et al. 2017; Pelkey et al. 2017). Double knockdown of *NR2F1* and *NR2F2* in the neurosphere and the developing mouse forebrain caused sustained neurogenesis and the prolonged generation of early-born neurons (Naka et al. 2008). Moreover, loss of NR2F1 caused an imbalance of excitatory/inhibitory neuron differentiation (overproduced GABAergic inhibitory interneurons and underproduced glutamatergic excitatory neurons) (Zhang et al. 2020), and altered percentages of different cortical interneuron subtypes (Lodato et al. 2011). All these indicate an underlying link between *MIR155* deletion-induced neuronal phenotypes and downregulation of *NR2F1* and *NR2F2*. In brain development, NR2F1 is crucial for establishing cortical patterning (Zhou et al. 1999; Zhou et al. 2001; Armentano et al. 2007) and hippocampal development (Yang et al. 2023) and NR2F2 plays a vital role in the development of the amygdala (Tang et al. 2012) and hippocampus (Yang et al. 2023). Our results further demonstrate that expressions of *NR2F1* and *NR2F2* are positively regulated by *MIR155*. Besides *miR155*, only three miRNAs have been implicated in positively regulating the expression of their target mRNAs under specific cellular conditions (Vasudevan et al. 2007; Place et al. 2008; Zhang et al. 2014). Thus, our studies and others underscore the importance of the *miR155*-NR2F1/NR2F2 pathway in the development and function of GABAergic interneurons.

Our findings that *miR155* deletion enhanced the generation of GABAergic interneurons in cortical organoids and, especially of PV-positive hippocampal GABAergic interneurons in *APP/PS1* mice are the first demonstration of a functional role for *miR155* in development and function of GABAergic interneurons. PV-positive interneurons control network synchrony and the generation of functional oscillations (Sohal et al. 2009; Hu et al. 2014) and their impairments can cause network abnormalities and memory deficits in AD patients and in mouse models of AD (Verret et al. 2012; Palop and Mucke 2016). Over the past decade, many studies have explored the restoration of PV interneuron function to address AD pathology and cognitive impairment in AD mouse models (Verret et al. 2012; Iaccarino et al. 2016; Palop and Mucke 2016; Martinez-Losa et al. 2018; Etter et al. 2019; Martorell et al. 2019). These studies have employed various approaches, including chemogenetics, optogenetics, genetic manipulations, and interneuron transplants. We reported that constitutive absence of *miR155* in *APP/PS1* mice prevented impaired learning behavior performance in the Barnes maze test, suggesting a beneficial effect of *miR155* deletion on learning behavior in the presence of early onset AD *APP* and *PSEN1* mutations and amyloidopathy (Readhead et al. 2018; Readhead et al. 2020). In the same study, field electrophysiology in hippocampal slices showed that *miR155* deletion partially rescued synaptic plasticity in 10-month-old *APP/PS1* mice but induced synaptic dysfunction in WT mice (Readhead et al. 2020). The present study supports the notion that using *miR155* inhibitors to improve PV interneuron function is a promising therapeutic strategy for network dysfunction in AD. This pathway may be particularly relevant to AD in DS patients, in whom *MIR155* upregulation is genetically programmed and defects in generation and maturation of GABAergic interneurons has been demonstrated in both differentiated hiPSCs derived from DS patients (Huo et al. 2018) and postmortem brain tissues from elderly DS patients (Ross et al. 1984; Kobayashi et al. 1990). Taken together, our findings significantly extend the classical paradigm for the cell-type-specific role of *miR155* in the pathogenesis of AD and other neurodegenerative diseases, in which *miR155* has, until now, been studied solely as a master regulator of microglia and neuroinflammation.

## Materials and methods

### Animals

The experimental procedures were conducted in accordance with NIH guidelines for animal research and were approved by the Institutional Animal Care and Use Committee (IACUC) at the Icahn School of Medicine at Mount Sinai. All mice were on a C57Bl6/J background. *APP^KM670/671NL^/PSEN1^Δexon9^* (= *APP/PS1*) (Jankowsky et al. 2004), and *miR155* knockout (*miR155^−/−^*) (Thai et al. 2007) mice were obtained from Jackson Laboratories. *APP/PS1* mice were crossed with *miR155^−/−^* mice to obtain WT, *miR155^+/−^*, *miR155^−/−^*, *APP/PS1*, *APP/PS1-miR155^+/−^*, and *APP/PS1-miR155^−/−^* mice. Male and female mice were used for immunohistochemistry, transcriptomic analyses, the quantification of target cell numbers, amyloid plaque positive area quantification and western blot analyses. If applicable, the use of both sexes is specified in the figure legends. 4- and 8-month-old mice were sacrificed by decapitation. One hemisphere was collected and immersion-fixed in 4% paraformaldehyde (PFA, w/v in PBS) for immunohistochemistry analysis. The other hemisphere was dissected, and the prefrontal cortex (PFC) and hippocampi were collected for qPCR and transcriptomic analysis. PFC, hippocampi, and cerebral hemispheres were snap-frozen and stored at -80 °C prior to RNA isolation or biochemistry analysis.

### Maintenance and culture of human induced pluripotent stem cells (hiPSCs)

All stem cell work was performed at the New York Stem Cell Foundation Research Institute (NYSCF). hiPSC cell lines, three AD lines (AD male (parental control), APOE3/3 background) AJ0083, AJ0094 and AJ0123 from ROSMAP cohorts (Bennett et al. 2018), were generated by NYSCF using a modified mRNA reprogramming method and were characterized for the typical hiPSC pluripotency, a normal karyotype and absence of mycoplasma contamination. hiPSCs were maintained in 6-well plates (Corning) in feeder-free conditions using Geltrex in complete mTeSR1 medium in a humidified incubator (5% CO^2^, 37°C). hiPSCs were fed fresh media daily and passaged using ReLeSR every 7-8 days.

### Generation of isogenic MIR155-deleted hiPSC lines

We designed a pair of sgRNAs (Supplemental Fig. S1B and Supplemental table S3) targeting exon 3 of *MIR155* host gene (*MIR155HG*) to remove a short (about 140 bp) genomic fragment containing pre-*MIR155* (65 bp) using the CRISPR Guide RNA design tool (https://www.benchling.com/crispr).

### Electroporation

hiPSCs with 70 to 80% confluence were dissociated by treating with Accutase and 10 µM ROCK inhibitor for 10 min. After spinning down hiPSCs at 300 r.c.f for 5 min, cells were counted, and 2 million cells were subjected to electroporation. Electroporation was performed using Nucleofector - Amaxa and Human Stem Cell Nucleofector Kit 1(Lonza) according to the manufacturer’s instructions. In brief, cells were resuspended in 100 µL of reaction buffer (82 µL Nucleofector solution and 18 µL Supplement) from the kit and 25 µL of RNPs (Ribonucleoprotein particles) and transferred to a Nucleocuvette. RNPs were complexes formed by combining 15 µL of sgRNA (100 µM, resolved in pH 7.0 TE buffer) and 10 µL of Cas9 (Alt-R S.p.Cas9 Nuclease V3 (Integrated DNA Technologies), 20 µM, resolved in pH 7.0 TE buffer) and followed by incubation at room temperature for 20 minutes. 2 µL of puromycin plasmid (0.5 µg/µL) were added to cell suspension. After nucleofection with the protocol B-016, cells were transferred to Geltrex-coated cell culture plates and cultured in complete mTeSR1 medium containing 10% CloneR for 2 days. Puromycin (1.0 µg/mL) was added to the medium for 1 day to select for electroporated hiPSC clones.

### PCR, DNA Electrophoresis and DNA Sanger sequencing

After 1-day puromycin selection, each colony was transferred to one well of a 48-well plate coated with Geltrex and maintained in complete mTeSR1 medium until the colony grew large enough to be transferred to a 6-well plate for further expansion. After the second transfer, hiPSCs in the original plate were dissociated, and genomic DNA was extracted. Genotyping primers (Supplemental Fig. S1B and Supplemental table S3) were used to amplify DNA fragment in *MIR155HG* gene and PCR products were subjected to DNA electrophoresis and submitted to GENEWIZ for Sanger sequencing.

### Karyotyping

To identify and evaluate the size, shape, and number of chromosomes in hiPSCs, we performed karyotyping (Giemsa banding and FISH (fluorescence *in situ* hybridization)) after Sanger sequencing. hiPSCs were cultured on Geltrex-coated 6-well plates in complete mTeSR1 media until 60% confluence and then sent to the Tumor CytoGenomics laboratory at the Icahn School of Medicine at Mount Sinai for karyotyping.

### Cortical neuron differentiation

hiPSC-derived cortical neurons were generated as previously described (Chambers et al. 2009; Qi et al. 2017) (Supplemental Fig. S2A). For neural differentiation to generate NSCs, hiPSCs were dissociated with Accutase and replated on 12-well tissue culture plates coated with Geltrex in complete mTeSR1 medium with 10 µM ROCK inhibitor. When cells were 100% confluent, the medium was replaced with Neural Induction medium (NIM, 50% DMEM/F12 medium, 50% Neurobasal medium, 2% B27 minus Vitamin A, 1% N2 supplement, 2 mM GlutaMAX and 100 U/mL Penicillin-Streptomycin) (day in vitro 0 (DIV0)) and maintained for 15 days. Inhibitors used in dual-SMAD inhibition protocol (LDN/SB/XAV, DIV0-10; XAV, DIV11-15) included LDN193189 (100 nM), SB431542 (10 µM) and XAV939 (1.0 µM). On DIV15 hiPSC-derived NSCs were dissociated using Accutase and can be stocked in Synth-a-Freeze™ Cryopreservation Medium (Thermo Fisher Scientific) in liquid Nitrogen tank.

For terminal differentiation to generate functional cortical neurons, hiPSC-derived NSCs (DIV15) were replated at 200,000-300,000 cells/cm^2^ in NIM medium with 10 µM ROCK inhibitor on dried poly-l-ornithine (Sigma-Aldrich) and laminin-coated 6-well plates or 8-well slide chambers. Cells were left to adhere overnight. On DIV16 NIM medium with ROCK inhibitor was replaced with Neuron Maturation media (BrainPhys™ Neuronal Medium, 2% B27 with Vitamin A, 2 mM GlutaMAX and 100 U/mL Penicillin-Streptomycin) supplemented with PD0325901 (10 µM), SU5402 (10 µM), DAPT (10 µM), BDNF (40 ng/mL), GDNF (40 ng/mL), Laminin (1.0 µg/mL), dBcAMP (250 µM) and L-Ascorbic acid (200 µM). During DIV16-23 Neuron Maturation media were half-changed every other day. After DIV24, the medium was replaced with Neuronal Maintenance media (BrainPhys™ Neuronal Medium, 2% B27 with Vitamin A, 2 mM GlutaMAX and 100 U/mL Penicillin-Streptomycin) supplemented with BDNF (40 ng/mL), GDNF (40 ng/mL), Laminin (1.0 µg/mL), dBcAMP (250 µM) and L-Ascorbic acid (200 µM). After DIV32, hiPSC-derived cortical neurons were active at this stage which can be confirmed with Ca^2+^ imaging. Between DIV50 and DIV60 hiPSC-derived cortical neurons were used for electrophysiological recordings.

### Electrophysiological recordings and analysis

For whole-cell recordings, cortical neurons derived from wild type and *MIR155*-deleted hiPSC lines between DIV50 and 60 were visualized using an upright Olympus BX51WI microscope equipped with a 40X objective and differential interference contrast optics. Cortical neurons were constantly perfused with 95% O^2^/5% CO^2^ in BrainPhys® medium preheated to 30-31°C. Patch electrodes were filled with internal solutions containing 130 mM K^+^gluconate, 6 mM KCl, 4 mM NaCl, 10mM Na^+^HEPES, 0.2 mM K^+^EGTA; 0.3mM GTP, 2mM Mg^2+^ATP, 0.2 mM cAMP, 10mM D-glucose. The pH and osmolarity of the internal solution were adjusted to resemble physiological conditions (pH 7.3, 290– 300 mOsmol). Current- and voltage-clamp recordings were carried out using a Multiclamp 700B amplifier (Molecular Devices), digitized with Digidata 1440A digitizer and fed to pClamp 10.0 software package (Molecular Devices). For spontaneous EPSC recordings, cortical neurons were held at a chloride reversal potential of -75 mV. Data processing and analysis were performed using ClampFit 10.0 (Molecular Devices) and MatLab (MathWorks) software. P-values to compare neurons alone to neurons with *MIR155* deletion were calculated using a two-tailed, unpaired t test.

### Isolation and characterization of lentivirus-mediated MIR155-overexpressing hiPSC-derived NSCs and GABAergic interneurons

Human *MIR155* (*hsa-MIR155*) precursors and approximately 100-bp upstream and downstream flanking genomic sequences were PCR amplified and cloned into a self-inactivated (SIN) lentiviral vector to generate pLV-miRNA vectors (Supplemental Fig. S2D). The cloning site of pre-miRNA genomic fragments is within the intron of human housekeeping gene EF1α promoter region. The rPuro gene product, expressed from the EF1α promoter, is the red fluorescent (mCherry) puromycin-N-acetyltransferase, and the transduced cells display red fluorescence at excitation/emission wavelengths of 587/610 nm. The miRNA lentiviral stock is prepared from cotransfecting HEK 293T cells with the pLV-miRNA plasmid and plasmids expressing Gag-Pol gene products and the vesicular stomatitis virus envelope G (VSV-G). The lentiviral supernatants were collected at 48 hours post transfection and stored at -70°C. The titer of the virus is generally above 1 X 10^7^ infectious units per mL (IU/mL). The miRNA of interest is delivered into cells by lentiviral transduction.

Dissociated hiPSC (cell line AJ0083)-derived NSCs were cultured on Geltrex-coated 6-well or 96-well plates in NIM medium containing 10 µM ROCK inhibitor. For lentivirus infection, we used lentivirus diluted in NIM medium containing 8 µg/mL polybrene (R&D Systems) to directly infect hiPSC-derived NSCs at a range of MOIs (1 to 20), which was calculated based on the starting virus concentration (2 X 10^7^ PFU/mL) and initial seeding density. To increase transduction efficiency, the plates were spun down at 1000 r.c.f. for 30 min at room temperature. After a 3-day infection, the medium was replaced with fresh NIM medium containing FGF2 (20 ng/mL), EGF (20 ng/mL) and puromycin (1.0 µg/mL) to select and expand for stably transduced cells. MiRNA qPCR analysis and double immunolabeling with mCherry and markers of NSCs (NESTIN, SOX2, SOX1 and Ki67) were performed for characterization of lentivirus-mediated *MIR155*-overexpressing hiPSC-derived NSCs (Supplemental Fig. S2E-F). All virus work was approved by the Mount Sinai Institutional Biosafety Committee.

hiPSC-derived cortical GABAergic interneurons were generated as described (Maroof et al. 2013) (Supplemental Fig. S2B). Briefly, dissociated hiPSC-derived NSCs (DIV15) were cultured for 8 days on Geltrex-coated 6-well plates in NIM medium containing 10 µM ROCK inhibitor, 1.0 µM purmorphamine and 5 nM SHH, a treatment condition referred to as “SHH” which enabled rapid and robust induction of ventral forebrain progenitor populations. For terminal differentiation to generate functional cortical GABAergic interneurons, hiPSC-derived NSCs (DIV23) were replated at 200,000-300,000 cells/cm^2^ in NIM medium with 10 µM ROCK inhibitor on dried poly-l-ornithine (Sigma-Aldrich) and laminin-coated 6-well plates or 8-well slide chambers. Cells were left to adhere overnight and on DIV24 NIM medium with ROCK inhibitor was replaced with Neuron Maturation media (BrainPhys™ Neuronal Medium, 2% B27 with Vitamin A, 2 mM GlutaMAX and 100 U/mL Penicillin-Streptomycin) supplemented with BDNF (40 ng/mL), GDNF (40 ng/mL), Laminin (1.0 µg/mL), dBcAMP (250 µM) and L-Ascorbic acid (200 µM). During DIV24-70 Neuron Maturation media were half-changed every other day.

### Cortical organoid differentiation

Cortical organoids were generated from hiPSCs as previously described, with modifications noted below (Madhavan et al. 2018) (Fig. 5A).

For patterning of neurocortical organoids, hiPSCs cultured on Geltrex were disassociated with Accutase to a single-cell suspension, to which was added an appropriate volume of mTeSR1 medium containing ROCK inhibitor (10 µM) to obtain 20,000 live cells per 200 µL. Then 200 µL of single-cell suspension were replated in each well of an ultralow-adherence V-bottom 96-well plates (S-Bio Prime; MS-9096VZ) and the plate was centrifuged at 1,000 rpm for 5 minutes at room temperature. The following day the medium was changed to mTeSR1 medium without ROCK inhibitor. On differentiation day 1, the cells were induced in 200 μL of Organoid Starter Media with 250 nM LDN193189 and 10 μM SB431542. Organoid Starter Media was DMEM/F12 containing 1% N2 supplement, Insulin (25 µg/mL), MEM non-essential amino acids (NEAA), Glutamax, β-mercaptoethanol, and 100 U/mL penicillin-streptomycin. The same media were used for the next 5 days with daily half-media changes, after which the media were changed to Organoid Differentiation Media. Organoid Differentiation Media was Neurobasal-A medium containing B-27 supplement without vitamin A, Glutamax, and 100 U/mL penicillin-streptomycin. From differentiation day 7 to day 25, 20 ng/mL FGF2 and 20 ng/mL EGF were added to Organoid Differentiation Media. Organoids were cultured in 96-well plates through day 20 and fed every other day with Organoid Differentiation Media. On day 20, organoids were transferred to ultralow-attachment 24-well plates (Corning; CLS3473) at a density of one organoid per well and cultured thus through the remainder of the protocol. Neural differentiation was induced between days 27 and 41 by supplementation of Organoid Differentiation Media with 20 ng/mL BDNF and 20 ng/mL NT3. Half-media changes were performed every other day between days 17 and 41.

To improve the maturation of cortical organoids, beginning on day 42, organoids were cultured in Organoid Maturation Media (Neurobasal-A medium containing B-27 supplement with vitamin A, Glutamax, and 100 U/mL penicillin-streptomycin) and 200 μM Vitamin C and 250 μM dbcAMP were added into the every-other-day media changes for two months. After day 100, mature cortical organoids were maintained in Organoid Maturation Media with every-other-day media changes until completion of the experiment.

### Immunocytochemistry and immunohistochemistry

Sixteen cortical organoids at day 10 of differentiation per batch of culture were pooled together for whole-mount staining and initially fixed with 4% ice-cold PFA for 1 hour at 4°C, washed three times in PBS and penetrated with PBS/0.3% Triton X-100. For immunocytochemistry (ICC), cells were washed three times with DPBS (1X) and fixed with cold 4% PFA for 15 min at 4°C, followed by three washes for 10 min each with PBS/0.1% Triton X-100 (PBST). The fixed cells or organoids were blocked with 10% goat or donkey serum/PBST for 1 hr at room temperature. The cells or organoids were then incubated with primary antibodies at the appropriate dilutions in 5% goat or donkey serum/PBST at 4°C overnight. The next day, cells or organoids were washed 3 times with PBST for 10 min each and stained with Alexa Fluor conjugated secondary antibodies (Thermo Fisher Scientific) at 1:400 for 2 hours at room temperature in the dark. After a final three washes with PBST for 10 min each and DAPI staining for 5 min, the slides with cultured cells or mounted organoids were coverslipped with ProLong Gold Antifade Mountant (Thermo Fisher Scientific).

Sixteen cortical organoids at day 42 or 66 of differentiation per batch of culture for immunohistochemistry were pooled together and initially fixed with 4% ice-cold PFA for 2 hours at 4°C, washed three times in PBS, and equilibrated with 30% sucrose at 4°C overnight. Then organoids were embedded in O.C.T (Tissue-Tek) and sectioned at 10 μm using a cryostat and the sections were mounted onto slides and stored at 4°C. For immunohistochemistry (IHC), mouse brains were collected and fixed in 4% PFA at 4°C overnight. Brains were then washed three times with PBS and placed in sucrose solution (30% w/v in PBS) overnight before being embedded in O.C.T (Tissue-Tek). Embedded tissue was sectioned at 30 μm using a cryostat and free-floating sections were stored in Cryopreservation medium at minus 20°C until staining. Sections were blocked in blocking solution (1X PBS, 0.3% Triton X-100 and 10% goat serum) for 1 hr at room temperature with gentle shaking. Prior to blocking, heat mediated antigen retrieval (The immunostaining experiments of organoid sections skipped this step) was performed by placing free-floating sections in a 1.5 mL micro centrifuge tube containing 1 mL of citrate buffer solution (pH 8.0) (Sigma) and placing in a pre-heated temperature block set at 80°C. Tissue was heated for 30 min at 80°C then removed and allowed to cool to room temperature before washing with PBS 3 times for 10 min each and then proceeding with blocking step. The sections were then incubated with primary antibodies at appropriate dilutions in blocking solution at 4°C overnight with slight shaking. The next day, sections were washed 3 times with PBST for 10 min each and stained with Alexa Fluor conjugated secondary antibodies at 1:400 for 2 hours at room temperature with slight shaking in the dark. After secondary staining, sections were washed in PBST 3 times for 10 min each and mounted on glass slides. After mounting, slides were coverslipped with DAPI-counterstain mounting media (Fluoromount, Southern Biotech).

Antibodies used in this study were verified by immunostaining mouse brain sections (data not shown) and are listed in Supplemental table S1. DAPI was used for counterstaining the nuclei. Stained sections were analyzed with Leica SP5 DMI confocal microscope (Leica) and Zeiss LSM980 Airyscan 2 confocal microscope (Zeiss). Images were assembled using Adobe Illustrator.

Mouse brain confocal images were thresholded and binarized using the Fiji ImageJ algorithm. The areas of the region of interest (ROI) in mouse hippocampi, DG, GCL, SGZ and hilus, were outlined by manually tracing with ImageJ software, and were calculated using the “measure” function. The numbers of RGL-NSCs (Sox1-, Sox2- and/or Gfap-positive cells) and GABAergic interneurons in ROIs were manually counted and numbers were normalized to the area of each ROI. Values were expressed as percentage of the control group.

### Western blotting

Aliquots of protein lysates (30 µg) prepared from mouse brains or hiPSC-derived cells and homogenized in RIPA buffer containing dissolved Protease Inhibitor Tablets (Thermo Fisher Scientific) were loaded in Criterion XT 4-12% Bis-Tris gels and transferred onto PVDF membrane (0.45 μm; Millipore, Billerica, MA, USA). Membranes were probed with primary antibodies (Supplemental table S1) at 4°C overnight with slight shaking. The next day, membranes were washed 3 times with TBST for 10 min each and subsequently incubated with anti-rabbit or anti-mouse HRP-conjugated secondary antibodies (1:2000, catalog #PI-1000 or PI-2000, Vector laboratories) at room temperature for 2 hours. After washing 3 times with TBST for 10 min each, membranes were developed with ECL Western blotting substrate (Pierce, Rockford, IL, USA) and were imaged on ChemiDoc MP imaging system (Bio-Rad). Normalization was achieved using GAPDH antibody (Supplemental table S1). The integrated density of immunoreactive bands was measured using Fiji software (ImageJ).

### RNA isolation, qPCR and miRNA qPCR analysis

Snap frozen mouse brain tissues from 4- and 8-month-old mice and hiPSC-derived cells were homogenized in QIAzol Lysis Reagent (Qiagen). Total RNA purification was performed with QIAGEN miRNeasy Mini Kit (Valencia, CA) following manufacturer’s guidelines. RNA quantification and quality were assayed using a NanoDrop 2000 (Thermo, Wilmington, DE). For reverse transcription, 1.0 μg of total RNA was transcribed into cDNA using a High-Capacity RNA-to-cDNA™ Kit (Applied Biosystems). Quantitative real-time PCR (qPCR) was performed in a StepOne Plus system (Applied Biosystem) using SYBR Green I Master (a hot start reaction mix, Roche, Germany) and data were normalized to the average of *GAPDH* and *HPRT1* expression. To reduce non-specific signals, primer sequences were designed using LightCycler Probe Design Software 2.0 (Roche, Germany) such that they spanned splice junctions, and they are listed in Supplemental table S2 and validated in pga.mgh.harvard.edu/primerbank/. Amplifications were performed in 20 μL containing 1.0 μL of each primer (5 μM), 10 μL SYBR Green I Master, 3 μL H_2_O and 5 μL of 20-fold diluted cDNA. Forty PCR cycles were performed with a temperature profile consisting of 95°C for 10 sec, 60°C for 15 sec and 72°C for 15 sec. The melting curve of each PCR product was determined to ensure that the observed fluorescent signals were only from specific PCR products, which were further verified by DNA Sanger sequencing. After each PCR cycle, the fluorescent signals were detected at 95°C for 5 sec and 65°C for 60 sec to melt primer dimers (Tms of all primer dimers used in this study were <76°C).

In this study, TaqMan™ Small RNA Assays were performed to detect and quantify mature miRNAs. For miRNA reverse transcription (RT), 15 ng of total RNA was transcribed into cDNA in RT Reaction Mix containing RT Primers (5X) (Supplemental table S2). MiRNA qPCR was performed in the StepOne Plus system using TaqMan® Gene Expression Master Mix containing TaqMan™ Small RNA Assay (20X) (Supplemental table S2) and 2-fold diluted cDNA templates. Data were normalized to expression of endogenous control *RNU49* (small nucleolar RNA, C/D box 49A, U49). The temperature profile of qPCR was described as above but the melting curve step was skipped during miRNA qPCR.

All qPCRs were performed on three biological replicates and were repeated using more than three independent samples, and data plotted as mean.

### miRNAscope

For miRNAscope (FISH) to detect *miR155* expression in hippocampal cells (NSCs and glutamatergic neurons and/or GABAergic interneurons), mouse brains were dissected and snap-frozen in liquid nitrogen before being embedded in O.C.T (Tissue-Tek). miRNAscope was conducted on mouse brain sections using RNAscope™ Plus smRNA-RNA HD Reagents Kit (Advanced Cell Diagnostics, catalog No. 322785) and miRNAscope™ Probe-SR-mmu-miR-155-5p-S1 (Advanced Cell Diagnostics, catalog No. 887751-S1) according to the manufacturer’s instructions with some modifications. The dehydrated tissue sections were treated with hydrogen peroxide for 10 min at room temperature. After rinsing with water, the sections were treated with protease III for 20 min at 40°C using the HybEZ hybridization system. After washing with water, the sections were incubated with the mmu-miR-155-5p-S1 probe for 2 hours at 40°C using the HybEZ hybridization system. After washing with RNAscope wash buffer, amplifiers 1 to 3 were used following the manufacturer’s protocol and the tissue sections were covered in HRP-S1 reagent and incubated for 15 min at 40°C. After washing with RNAscope wash buffer, the sections were covered with diluted fluorophore Opal 570 and incubated for 30 min at 40°C. After washing with RNAscope wash buffer, the sections were covered with HRP Blocker and incubated for 15 min at 40°C and followed by RNAscope wash buffer. Finally, slides were coverslipped with DAPI-counterstain mounting media (Fluoromount, Southern Biotech).

### RNA sequencing data analysis

RNA isolation was performed on the lysates of WT, *MIR155*KO Clone 5 and *MIR155*KO Clone 25 hiPSC-derived cells, and *MIR155*-overexpressing NSCs and cortical neurons, according to the manufacturer’s protocol (miRNeasy mini-kit, Qiagen). Isolated RNA samples were submitted to Novogene Corporation Inc. for analysis of RNA quality and integrity. All samples passed quality control, and RNA-seq libraries were prepared for sequencing using NovaSeq (Supplemental tables S4-5). Paired-end RNA sequencing FASTQ files for all samples were aligned to Homo sapiens Reference genome (Grch38) using STAR read aligner and mapped reads were summarized to gene counts using subread function of FeatureCounts. Genes with expression (< 1 FPKM) across all samples were filtered from all subsequent analysis. Hierarchical clustering and principal component analysis (PCA) were preformed as part of quality control to visualize biological variance between replicates, which were further exhibited by Volcano plots and Venn diagrams. We then performed differential gene expression analysis to compare all cell types with each other on normalized counts with EdgeR. Multiple biological replicates were used for all comparative analysis. Differentially expressed genes (DEGs) were identified at a false discovery rate (FDR) of 0.05. To identify biological pathways that may be differentially dysregulated by *MIR155*KO or *MIR155* overexpression, we performed gene set enrichment analysis (GSEA) on the identified DEGs by using Enrichr, GSEA-MSIGDB and IPA (Ingenuity Pathways Analysis), where we identified enriched biological pathways, gene ontology sets, and mRNA targets of miRNA at an adjusted p-value < 0.05. In heatmaps (Fig. 3B, 3C, 3F, 3G, Supplemental Fig. S4A, S4C, S4E, S8I, S8J, S8P), X-axis (Horizontal) represents different comparative analysis groups (*MIR155*KO Clone 5 hiPSCs-derived cells vs WT hiPSCs-derived cells, *MIR155*KO Clone 25 hiPSCs-derived cells vs WT hiPSCs-derived cells, and *MIR155*-overexpressing cells vs Control cells). Y-axis (Vertical) represents genes. The top-right scale bars for color coding display the range of Log_2_ FoldChange that values greater than 0 indicate upregulation (red) and values less than 0 indicate downregulation (green).

### ATAC-sequencing, H3K4me3 and H3K27ac ChIP–sequencing and Hi-C sequencing data analysis

External validation datasets (Hu et al. 2021; Bendl et al. 2022; Girdhar et al. 2022; Rahman et al. 2023): MSBB RNA-seq of postmortem brains (Synapse ID: syn3157743), ATAC-seq on FANS (fluorescence-activated nuclear sorting)-sorted NeuN+/− nuclei from postmortem adult brains (Synapse ID:syn21531502), H3K4me3 and H3K27ac ChIP–seq of neuronal nuclei isolated from adult control brains (Synapse ID: syn25705564), and Hi-C sequencing on human fetal cortical plate and on FANS-sorted neuronal (NeuN+) and glial (NeuN-) nuclei isolated from adult prefrontal cortex (Synapse ID: syn26256090 and syn21760712).

ATAC-seq libraries were generated from neuronal and nonneuronal nuclei isolated by FANS from frozen postmortem human brain tissue dissections as described (Bendl et al. 2022). All libraries were processed by established computational pipeline that performs read mapping (STAR), peak calling (MACS), genotype calling (GATK and KING) and quality control checking. Extensive quality control of ATAC-seq libraries based on the cell type, sex and genotype concordance, as well as sample quality metrics and sequencing depth, yielded a total of 636 samples constituting a total of 19.6 billion read pairs with an average of 30.8 million nonduplicated read pairs per library.

The Hi-C sequencing data analysis was performed using a standard protocol (Hu et al. 2021; Bendl et al. 2022; Rahman et al. 2023). Hi-C sequencing data were aligned using the HiC-Pro strategy. Briefly, paired-end reads were mapped independently to human genome hg38 using bowtie2 in stringent mode with parameters “--very-sensitive -L 20 --score-min L,-0.6,-0.2 --end-to-end”. Then the chimeric reads that failed to align were trimmed after ligation sites (MboI “GATCGATC”) and mapped to the genome. All the aligned reads from two ends were then merged based on reads name and mapped to MboI restriction fragments using the hiclib package. After that, self-circles, dangling ends, PCR duplicates, and genome assembly errors were discarded. Samples of the same cell type were merged. We binned the interaction matrix at different resolutions and corrected it with iterative correction (ICE) for downstream analysis. Chromatin loops were called with HICCUPS for two different cell types independently. First, we converted the filtered interaction files into Juicer format with JuicerTools. Chromatin loops were called using juicer HICCUPS with bin sizes iterated from 10kb to 25kb by 1kb intervals and parameters “-k VC_SQRT -p 1 -i 3”. Only reproducible loops were retained and the highest resolution of the overlapping loops were used. Topologically associated domains (TADs) were identified with TopDom (v0.0.2(18)) at 10K resolution and 200kb window size. For compartment analysis, first, we calculated the genome-wide correlation matrix at 200Kb resolution with only intrachromosomal interactions. Then the first eigenvector of the correlation matrix was obtained and corrected the sign to have a positive correlation with GC content and gene density. The signs of the eigenvector were used to assign the genome into compartments A and B.

### Statistical analysis

No statistical methods were used to predetermine sample size and the experiments were not randomized. Experimental data were analyzed for significance using Excel. P<0.05 was considered statistically significant. All experiments were analyzed by ANOVAs followed by Tukey’s test, Dunnett’s test or unpaired Student’s t tests.

## Data availability

The GEO (Gene Expression Omnibus) database accession numbers for the RNA sequencing data reported in this paper are GSE277199 and GSE277200.

## Competing interest statement

Dr. Sam Gandy is a co-founder of Recuerdo Pharmaceuticals. He has served as a consultant in the past for J&J, Diagenic, and Pfizer, and he currently consults for Cognito Therapeutics, GLG Group, SVB Securities, Guidepoint, Third Bridge, MEDACORP, Altpep, Vigil Neurosciences, Eisai, Memory Garden, the Bell Law Firm, Alzheon, and Rylands Garth Legal Services. He has received research support in the past from Warner-Lambert, Pfizer, Baxter, and Avid.

## Acknowledgements

1. S. N., V. F., S. G. and M. E. E. acknowledge the support of NIH grant R01AG061894. S. G. acknowledges the support of NIH grant P30 AG066514 to Mary Sano. S. G. and M. E. E. acknowledge the support of NIH grants U01AG046170 and RF1AG058469 and the Cure Alzheimer’s Fund. This work was also supported by the New York Stem Cell Foundation Research Institute (NYSCF). We thank Tumor CytoGenomics laboratory at the Icahn School of Medicine at Mount Sinai for karyotyping, and the Microscopy Core for technical support for confocal imaging. We thank Novogene Corporation for performing RNA sequencing and the computational analysis. We thank Allen Pan for miRNAscope assay, Darlinda Shillingford for preparation of mouse brain sections, and members of the Ehrlich/Gandy lab and the mouse facility at the Icahn School of Medicine at Mount Sinai for the maintenance of mouse lines. ROSMAP is supported by P30AG10161, P30AG72975, R01AG15819, R01AG17917, U01AG46152, U01AG61356. ROSMAP resources can be requested at https://www.radc.rush.edu.

## Author contributions

X. D. Z. and M. E. E. conceived the project, designed experiments, and developed experimental protocols, tools, and reagents. V. F., S. N., S. G. and M. E. E. supervised the study. J. V. H. M. and A. A. prepared the mouse brain sections and performed immunostaining. M. B. performed components of the computational analysis. P. F. D. performed ATAC-sequencing and Hi-C sequencing data analysis. I. K. carried out electrophysiological recordings and analysis. X. D. Z. performed all the remained experiments and computational analyses. X. D. Z. wrote the first draft of the manuscript. J. V. H. M., V. F., S. N., S. G. and M. E. E. reviewed and edited the manuscript. All authors read and approved the paper.

